# Mapping multiregional image-localized biopsies to MRI habitats reveals biologically significant glioma tissue states and patterns of cellular subpopulations

**DOI:** 10.1101/2022.03.23.485500

**Authors:** Lee Curtin, Kamila M. Bond, Pamela R. Jackson, Andrea Hawkins-Daarud, Sebastian Velez, Neha Shakir, Javier C. Urcuyo, Gustavo De Leon, Kyle W. Singleton, Jazlynn M. Langworthy, Kamala R. Clark-Swanson, Yvette Lassiter-Morris, Lisa Paulson, Christopher Sereduk, Chandan Krishna, Richard S. Zimmerman, Devi Prasad Patra, Bernard R. Bendok, Kris Smith, Peter Nakaji, Kliment Donev, Leslie Baxter, Nhan L. Tran, Leland S. Hu, Christopher Plaisier, Maciej M. Mrugala, Alexander R.A. Anderson, Jack Grinband, Osama Al-Dalahmah, Joshua B. Rubin, Peter Canoll, Kristin R. Swanson

**Author notes:** Corresponding and lead author **Lead Contact:** Kristin R. Swanson, PhD, Cedars-Sinai, Advanced Health Science Pavillion, 127 S. San Vicente Blvd. 6th Floor, A6317A, Los Angeles, CA 90048. Co-first authors contributed equally (Lee Curtin, Kamila M. Bond).

## Abstract

**Background:** Magnetic Resonance Imaging (MRI) is the mainstay for neurosurgical oncology but not for informing us about glioma biology. An obstacle to developing MR-based glioma biomarkers is the absence of rigorous correlation between MRI features and glioma biology as assessed in multi-regional biopsies, within and across patients.

**Methods:** We directly addressed this obstacle by collating a unique cohort of 202 MRI-localized biopsies from 58 patients. We define a low-dimensional transcriptional pseudotime continuum along which heterogeneous high-grade glioma (HGG) samples organize both within and across patients.

**Results:** We observe three polarized transcriptional tissue states: infiltrated brain, immune/inflammatory, and proliferative associated with patterns of cohabitation of cellular subpopulations. The states and deconvolved populations show correlation with enhancement status on T1Gd MRI. Moreover, discrete MRI habitats, regions sharing common imaging features, defined as combinations of high or low signal intensity across multiparametric MRI revealed 14 MRI habitats. We order the MRI habitats according to the average pseudotime on the transcriptional continuum. We find that MRI habitats with low pseudotime (associated with early tumor development and diffusely invaded brain tissue) localized at the periphery of the tumor whilst high pseudotime either proliferative or immune/inflammatory states were towards the core of the lesion. We find that composition of MRI habitats is impacted by MGMT status.

**Conclusion:** This suggests that ongoing aggregation of MRI-localized biopsies may augment our projection of biology onto MRI habitats to support the noninvasive identification of cellular ecologies within and across each patient’s tumor.

**Lay Summary:** In notoriously heterogeneous and aggressive high-grade gliomas, even within a patient, different molecular patterns are present depending on where tissue is collected. Here, we collected multiple biopsies from each patient and recorded their locations on matched MRI. For each biopsy, we quantified the tumor, immune, and normal cell subpopulations present, detected patterns of cellular communities, and connected these cellular communities to MRI-defined regions, which allowed us to order MRI regions on a potential aggressiveness axis. This approach could allow for the tracking of the biological response of cellular communities to treatment in patients in real time from imaging alone.

**Key Points:**

- Trajectory inference of HGG samples reveals a continuum connecting three polarized tissue states reflecting transitions in cellular population ecologies and gene set enrichment
- Combining multiparametric MRI features into regional MRI habitats reveal clinically-distinct patterns of cell co-habitation and tissue states
- MGMT methylation status shifts the cellular patterns of cohabitation within MRI habitats

**Importance of the Study:** Regional heterogeneity is a defining characteristic of high-grade glioma (HGG). Mapping the transcriptomic biology of multiregionally sampled MRI-localized biopsies from high-grade gliomas onto a low-dimensional space reveals a continuum between three primary tissue states: infiltrated brain, immune/inflammatory, and proliferative. Each state is associated with distinct cell cohabitation ecologies. Transitions between the tissue states coordinate with changes in tumor composition, revealing potential direct connections between tumor biology shifts and their manifestation on clinical imaging.

## Introduction

High grade glioma (HGG) is genetically, transcriptionally, and compositionally heterogeneous within and between patients^1,2^. Even cells within the same biopsy can have different therapeutic sensitivities. Access to brain tumor tissue is limited, leading to an urgent clinical need for tools that can incorporate tumor heterogeneity to assess and/or predict treatment response in a non-invasive manner. Many attempts to characterize this heterogeneity have discretized samples into biological groups characterizing tumor cell populations rather than non-tumor cells.^3–9^ These models fail to embrace the evidence that glioma cells engage in reciprocal cross-talk with normal cells in the microenvironment^10–12^. Under this conceptual framework, the biology of tumor tissue is a direct reflection of its cellular constituents and the environmental variables that influence them. Work by Gill, Al Dalahmah and others has characterized these interactions into three primary tissue states^8,9^.

MRI is the mainstay of treatment assessment for HGG, but the mapping between signal intensities and the underlying biology are opaque. The prevailing gold standard for the radiographic evaluation of HGG is determined by the changes in the contrast-enhancing (CE) on T1-weighted MRI and/or non-enhancing (NE) tumor volumes on T2/FLAIR MRI considering patients’ steroid usage and clinical status^13^. This paradigm is fundamentally limited by the non-specificity of contrast enhancement on T1-weighted MRI and T2/FLAIR MRI hyperintensity. This tenuous relationship between MRI signal and biology is evident with the use of anti-angiogenic agents (i.e., bevacizumab), where an acute reduction in capillary permeability leads to a decrease in intratumoral contrast enhancement without a change in tumor burden^14^. Conversely, about a third of patients demonstrate an increase in CE and progressive NE extent upon the initiation of radiochemotherapy in a phenomenon called pseudoprogression^14,15^. It is assumed that T2/FLAIR hyperintensity represents infiltrative tumor, but vasogenic edema and reactive changes can also contribute to an abnormal T2/FLAIR signal in the peritumoral tissue. Metrics from advanced MRI modalities dynamic susceptibility contrast (DSC) and diffusion-weighted imaging (DWI) have shown promise in distinguishing progression from pseudoprogression^16^, and recent work has used these alongside standard imaging to distinguish tissue states in a recurrent setting^17^.

Early work established that multiparametric MRI could partition tumors into “habitats,” but these regions were largely inferential, lacking direct spatial biological validation^18–22^. Subsequent radiogenomic studies demonstrated that imaging features could predict molecular alterations such as IDH status, yet typically relied on single or non-spatially matched biopsies, overlooking the known spatial heterogeneity of the disease^23–27^. Other studies have emphasized voxel-wise predictions, alongside a growing recognition that rigorous validation requires spatially co-registered, multiregional tissue sampling^28–30^. In this context, our study proposes to incorporate 202 MRI-localized biopsies with matched bulk transcriptomics to provide ground truth validation of the biology underlying distinct MRI habitats. This addresses the longstanding MRI-biology-mapping validation gap, moving us beyond correlative radiogenomics toward true spatio-genomic mapping.

In this study, we leverage a manifold learning algorithm to organize a novel cohort of 202 MRI-localized HGG biopsies along transcriptional trajectories. This is supplemented with a deconvolution algorithm and gene signature enrichment to determine the cellular compositions and hallmark signatures associated with each biopsy. We connect the resultant spectrum of tumor ecologies with MRI habitats. The workflow (Figure 1) includes: 1) Segmentation of abnormal regions of T1Gd and T2 MRI, normalization and registration across all imaging protocols; 2) Monocle trajectory inference analysis to characterize transcriptional states and order samples by pseudotime; 3) CIBERSORTx deconvolution of each sample into 18 cell states, including malignant, immune and other; 4) Characterization of advanced MRI habitats based on binary “high” and “low” group combinations; 5) Comparison across imaging and transcriptional state/pseudotime/population outputs to determine population cohabitation patterns and their MRI habitat presentation.

**Figure 1.**
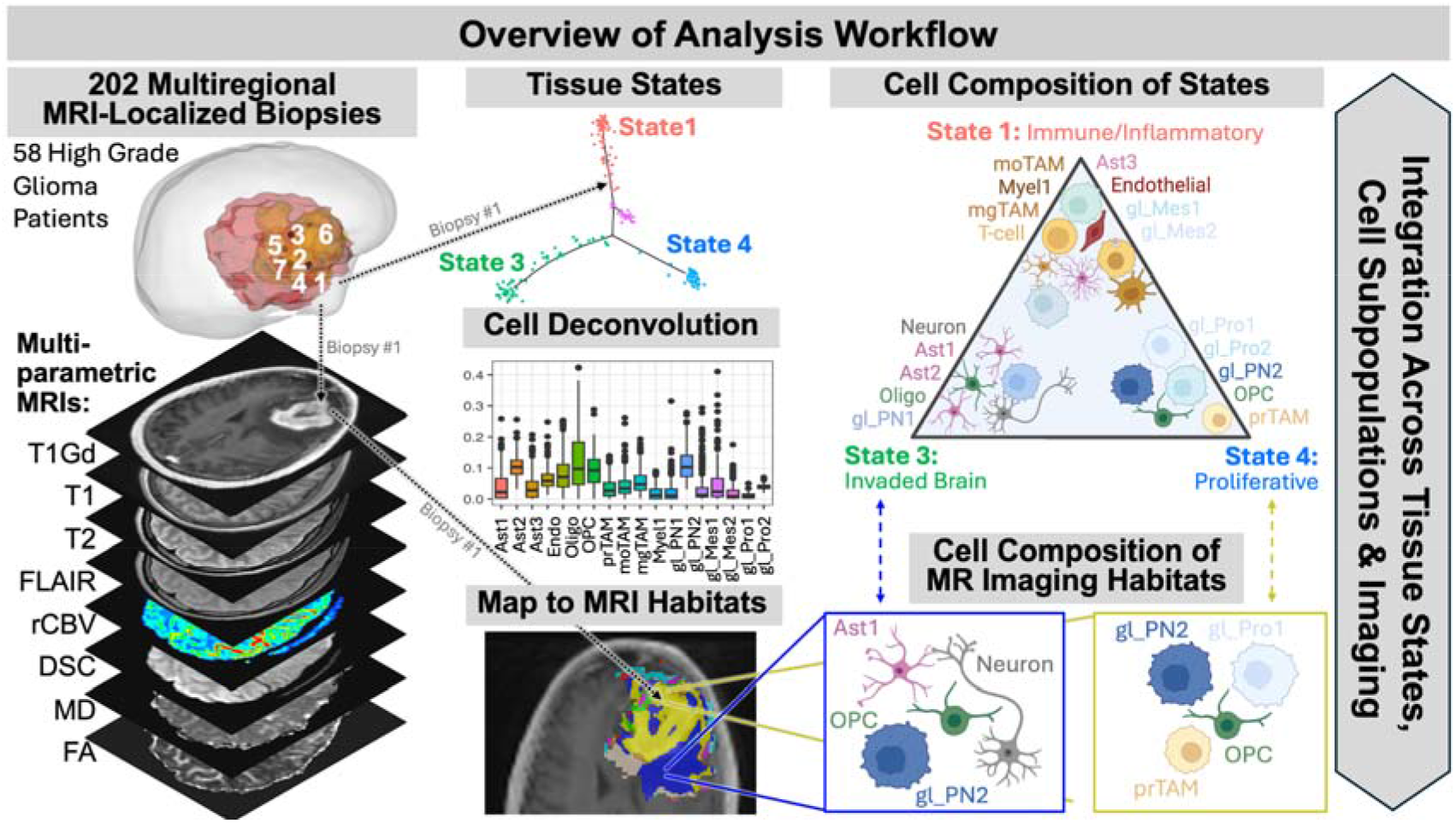
Overall workflow schematic. Bulk RNAseq from 202 multiregional biopsies from 58 glioma patients are distributed across a 3-armed tissue state graph. Each biopsy is deconvolved to allow connection of tissue state with cellular composition. Each biopsy location is also mapped to MRI. Multiparametric MRI allows creation of imaging distinct habitats as combinations of high and low signals from each MRI sequence. By aligning the multiregional biopsy samples to each MRI location, each MRI habitat can be characterized in terms of tissue state and cellular composition. This allows a direct linking of MRI changes with tumor biological shifts *in vivo* within each patient.

## Methods

### Tissue samples

*Image-Localized Biopsy Data*: After obtaining informed consent, we enrolled 58 patients (22F, 36M) with clinically-suspected high-grade glioma in our IRB-approved study. Patients underwent surgical tumor resection with intraoperative navigation, and a total of 202 biopsies were harvested, with 195 of those image localized.

### Deconvolving cell populations from RNA sequencing

We leveraged a snRNAseq data set^9^, derived from a combination of patients with low- and high-grade glioma as well as epilepsy, which contained eighteen cell states: six malignant, five immune, and seven “other” summarized in Table S1. We used CIBERSORTx^31^ to deconvolve each bulk RNASeq sample into relative abundances of each of these single-cell-defined populations and averaged across multiple runs (Figure S1).

### Trajectory inference and pseudotime ordering of TCGA microarray and image-localized RNAseq samples

We repurposed trajectory infererence^32^ often used for ordering single-cell RNAseq data to determine the relative ordering of our bulk biopsy data^33^. Here we used TCGA microarray and our biopsy RNAseq as inputs to Monocle to visualize the natural transcriptional organization of HGG. States for our image-localized biopsy data are presented as assigned by Monocle.

### Differential Gene Expression and Gene Set Enrichment Analysis

Differential expression used the “limma” package in R with a patient block design to account for multiple samples per patient^34^. The “fgsea” package was then used to perform gene set enrichment analysis (GSEA), with the hallmarks pathways from the MSigDB database as the reference^35^. Single sample GSEA was used to determine the pathway expression of each sample using the “gsva” package in R^36^. HGNCHelper was used for alignment between the image-localized and TCGA datasets^37^ (See Supplementary Methods).

### *MGMT* Methylation Inference

A surrogate of MGMT methylation was inferred for samples without clinical status, using a threshold based on the 75 of 202 samples with values (See Supplementary Methods). These inferred values were used only to supplement visualization of MGMT and not in statistical analyses due to variability in MGMT methylation and transcriptional expression present within tumors^38^.

### Statistical analyses

Patient IDs were included as a random effect in each test within “lmerTest” R package^39^, with Satterthwaite’s method used for comparisons of 2 groups, and anova tests implemented for comparisons with 3+ groups. A Benjamani-Hochberg correction was used when adjusting for multiple comparisons.

## Results

### Trajectory inference of HGG samples reveals three polarized tissue states

Bulk transcriptomic sequencing of 202 biopsies from 58 patients (mean=3.5 per patient), with clinically-suspected high-grade glioma underwent manifold learning algorithm trajectory inference using Monocle^40^. This origin (pseudotime = 0) was set to the end of the trajectory associated with non-neoplastic brain cells (Figure 2A). Patient demographics and biopsy characteristics are summarized in Tables S2 and S3, respectively. Monocle classified these into 5 states (Figure 2B, Table S4). State 2 was not assigned to any samples, so is excluded from analysis. Glioma tissue State 3 is to the left, while State 4 is on the bottom arm and s 1 and 5 are on the top arm. Bulk GSEA results and ssGSEA results show significant enrichment between states (Figures S2A and S2B). Top arm State 1 is associated with immune hallmarks. State 4 is associated with both proliferative populations and proliferation-related hallmarks. State 1 shows the most significant upregulation of hallmark signatures when compared against all other states, whereas State 3 shows the most significant downregulation. There is an insignificant bias for male samples towards higher pseudotime values (p=0.28) (Figure 2C). IDH mutant samples tend to occur in State 5 (Figure S3A, p=0.0122). MGMT methylated samples are more likely to be found in proliferative glioma State 4 than in immune/inflammatory glioma State 1 (Figure S3B, p=0.0449). TCGA sample classification overlaid on the trajectory (Figure S4A) confirms that the Monocle graph can significantly distinguish transcriptional states (X^2^=507, p<0.0001; Figure S4B).

**Figure 2.**
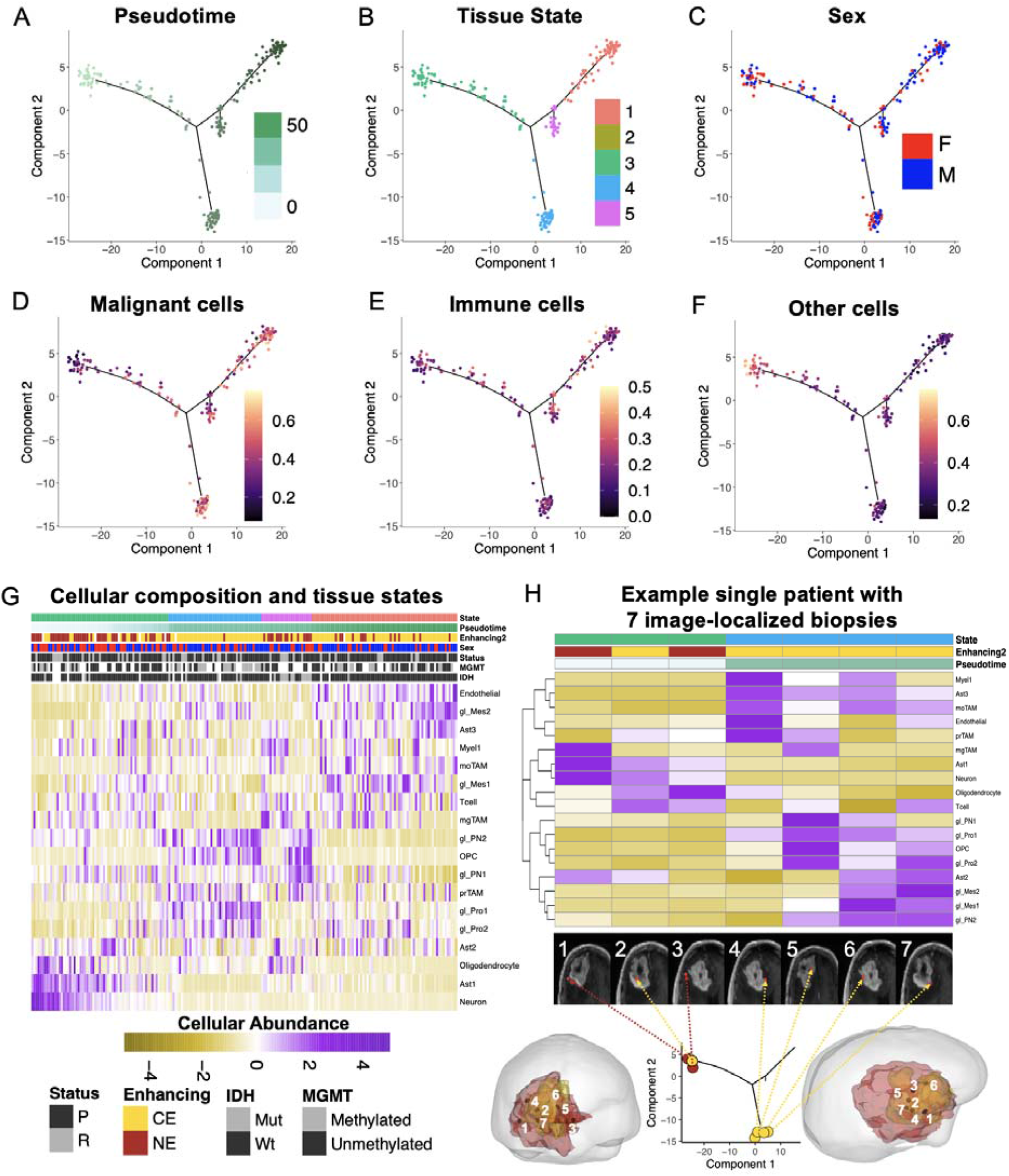
Trajectory inference of 202 HGG biopsies (195 image-localized) from 58 patients recapitulates the three arms, each of which has distinct cellular cohabitation patterns. **(A)** Pseudotime trajectory and **(B)** States derived from biopsies are similar to the 3 arms seen in TCGA (Figure S4A). **(C)** There is a sex bias in our samples such that female samples have lower pseudotime. **(D)** CIBERSORTx deconvolution of bulk RNAseq allows for the estimation of cell population abundances. Malignant cells are present on all arms of the trajectory, with overrepresentation in State 4, particularly proliferative glioma and one proneural glioma subpopulation (gl_PN2). **(E)** Immune cells (i.e., moTAM, Myel1, Tcell) are most abundant in State 1 with mesenchymal tumor, top arm. **(F)** Other cells (i.e., neurons, oligodendrocytes, astrocytes) are most populous in State 3, left arm. **(G)** Heatmap of relative cohabitation tendencies of cell populations ordered by Monocle state, then pseudotime. **(H)** Exemplar Patient 33 (59-year-old female with primary IDHwt GBM) with seven image-localized biopsies.

### States reflect transitions in cellular population ecologies and gene set enrichment

Overlaying CIBERSORTx-predicted cell state proportions onto the Monocle trajectory reveals population ecologies (Figure 2D,E,F; Table S1). The sums of malignant cells, immune cells, and other cells are highest on end States 4, 1 and 3 respectively (Figure 2D). Of the 18 different cell types, 16 of them showed significant differences between the 4 main Monocle-predicted states (Figure S5). Shannon indices were different between states (Figure S6). Transitions between states correlate with changes in composition (Figure 2G), with State 4 showing a mixture of populations consistent with a transitional state as seen on the Monocle trajectory (Figure 2G). HGG samples with low pseudotime are associated with early tumor development within State 3 (left arm) with a small amount of tumor cells (mostly proneural OPC-like gl_PN1) with a high fractional abundance of neurons, oligodendrocytes, and normal (non-reactive/protoplasmic) astrocytes (Ast1). As pseudotime increases to bottom arm State 4 there is a phased enrichment in OPCs and vascular-associated prTAMs coordinated with an increase in another proneural (NPC-like) glioma phenotype (gl_PN2) and proliferative glioma populations (gl_Pro1, and gl_Pro2). As pseudotime increases along the top arm, we find a small State 5 with enrichment in OPCs, plasticity/tumor–glial hybridization reactive astrocytes (Ast2), microglial-derived mgTAMs, inflammatory/metabolic stress myeloid cells (Myel1), as well as gl_PN1 tumor (with an associated increase in IDH1mut status - Figure S3A). Further along the top arm, State 1 mesenchymal immune/inflammatory tumor is enriched in endothelial cells, inflammatory astrocytes Ast3, gl_Mes1, gl_Mes2, and some T-cells.

### Mesenchymal immune/inflammatory tissue state 1 is associated with poor outcomes in female patients

Applying image-localized tissue state signatures to 525 TCGA patients as outlined in the Methods, near significant survival differences are detected between the two primary tumor-rich states, the mesenchymal immune/inflammatory State 1 (270 patients) and more proliferative glioma State 4 (62 patients, p=0.097, log-rank test, Figure S7A). Within female samples, State 4 (98 patients) had a significantly longer survival than State 1 (25 patients, p=0.04, Figure S7B), while no significant result was observed amongst male samples (p=0.65, Figure S7C).

### Cellular composition of samples within the same patient is highly variable

Figure 2H shows a case example of a 59-year-old female patient with newly diagnosed right frontal IDH wild-type, MGMT methylated GBM, who had eight image-localized biopsies collected (Figure 2H; 5CE, 2NE, 1 uncharacterized) and lived 174 days from diagnosis. The patient’s biopsies were distributed across the pseudotime trajectory and 2 states. Biopsies with similar imaging appearance (i.e., both NE or both CE) were associated with a spectrum of cellular composition similarity (Figure 2G, Figure S8). For example, samples 2 and 3 from this patient have nearly identical cell compositions (R=0.96), but one is CE and one is NE. Conversely, samples 2 and 7 are compositionally dissimilar (R=-0.14) yet they are both CE.

### Contrast-enhancing MRI regions are enriched in malignant cells

NE samples had a significantly lower pseudotime (p<0.0001). NE samples are more commonly found in States 3 and 5, while CE samples are found in States 1 and 4. (Figure 2G). Further, CE samples taken from the necrotic edge of the tumors were all observed to have high pseudotime values (Figure S9). Treatment status (i.e., primary and recurrent) has no clear relationship to states (Figure S10). Several cell types showed significant differences in abundance between CE and NE regions (Figure S11). We observed a higher relative abundance of malignant cells in CE than NE regions across all samples (p=0.0006), a trend not driven by primary/recurrent status nor patient sex (maintained significance). Amongst NE samples, there are significantly fewer malignant cells in recurrent biopsies compared to untreated primary biopsies (p=0.005), a result not seen amongst CE samples or all samples (p=0.58, p=0.72, respectively).

### NE regions are enriched in some normal brain cell populations

We observe more neurons in NE samples (p<0.0001; Figure S11), the significance of which is retained in both male and female samples (p=0.013 and p=0.0033, respectively), and primary and recurrent settings (p=0.0016 and p=0.0023, respectively). Protoplasmic astrocytes, Ast1, were more prevalent in NE samples (p<0.0001, Figure 2H) which appears more strongly in females than males, despite significance in each sex (p<0.0001 females, p=0.012 males). Lastly, normal oligodendrocytes are more common in NE samples (p=0.012). OPCs were not significantly different between CE and NE regions (p=0.78).

### Mesenchymal glioma cells are more commonly found in CE regions

There was an overrepresentation of gl_Mes2 and gl_Pro1 cells in CE compared to NE biopsies (p<0.0001, p=0.0087, respectively; Figure S11). Conversely, NE biopsies have more gl_PN1 cells than their CE counterparts (p=0.0070; Figure S11). While gl_Mes2 was significant regardless of sex (M: p=0.013, F: p=0.0019), females drove significance for gl_Pro1 (M: p=0.31, F: p=0.020) and gl_PN1 (M: p=0.31, F: p=0.0022). Reactive astrocytes, Ast3, also showed a small but significant enrichment in CE regions (p=0.018). There were no significant regional differences in gl_Mes1, gl_Pro2 or gl_PN2 across all samples or within patient sex.

### Different tumor associated macrophage populations are enriched in CE vs NE

Monocyte-like tumor associated macrophages, moTAMs, were more common amongst immune cells in CE whereas microglia-like mgTAMs were more common in NE samples (p=0.0087, p=0.024, respectively). When broken down by sex, the only signal that was preserved was mgTAM in female samples when adjusting for multiple comparisons (p=0.043).

### Endothelial cell abundance is correlated with contrast-enhancement

This is consistent with CE representing leaky angiogenic neovasculature allowing leakage of gadolinium^41^, the contrast agent used in MRI (p=0.00054). Endothelial cell abundance was found to significantly vary with Monocle states (p<0.0001) and was highest in State 1.

### MGMT methylation status is associated with shifts in cellular populations present with CE and NE

Consistent with the evolutionary concept that there is a cost to resistance^42^, in the competitive microenvironment CE core of the tumor, “responsive” methylated glioma samples have improved fitness showing an increased enrichment in actively proliferating glioma subpopulation, gl_Pro1 (p=0.0066), and moderately proliferative gl_PN2 (p=0.039). Conversely, the unmethylated samples are associated with a more mesenchymal glioma cell phenotype that has less proliferative potential (gl_Mes1, p=0.043). Additionally, amongst non-glioma, non-immune cells, OPCs dominate the CE TME for methylated samples (p=0.04).

### DTI’s mean diffusivity correlates with glioma cell abundance

In NE samples, the DTI metric apparent diffusion coefficient (ADC) has previously been shown to correlate with total cellularity of the sample^43^, not necessarily tumor cellularity. We ask the refined question if the relative abundance of glioma cells within the sample is reflected by ADC from DTI but the relationship did not reach significance (Figure 3D).

**Figure 3.**
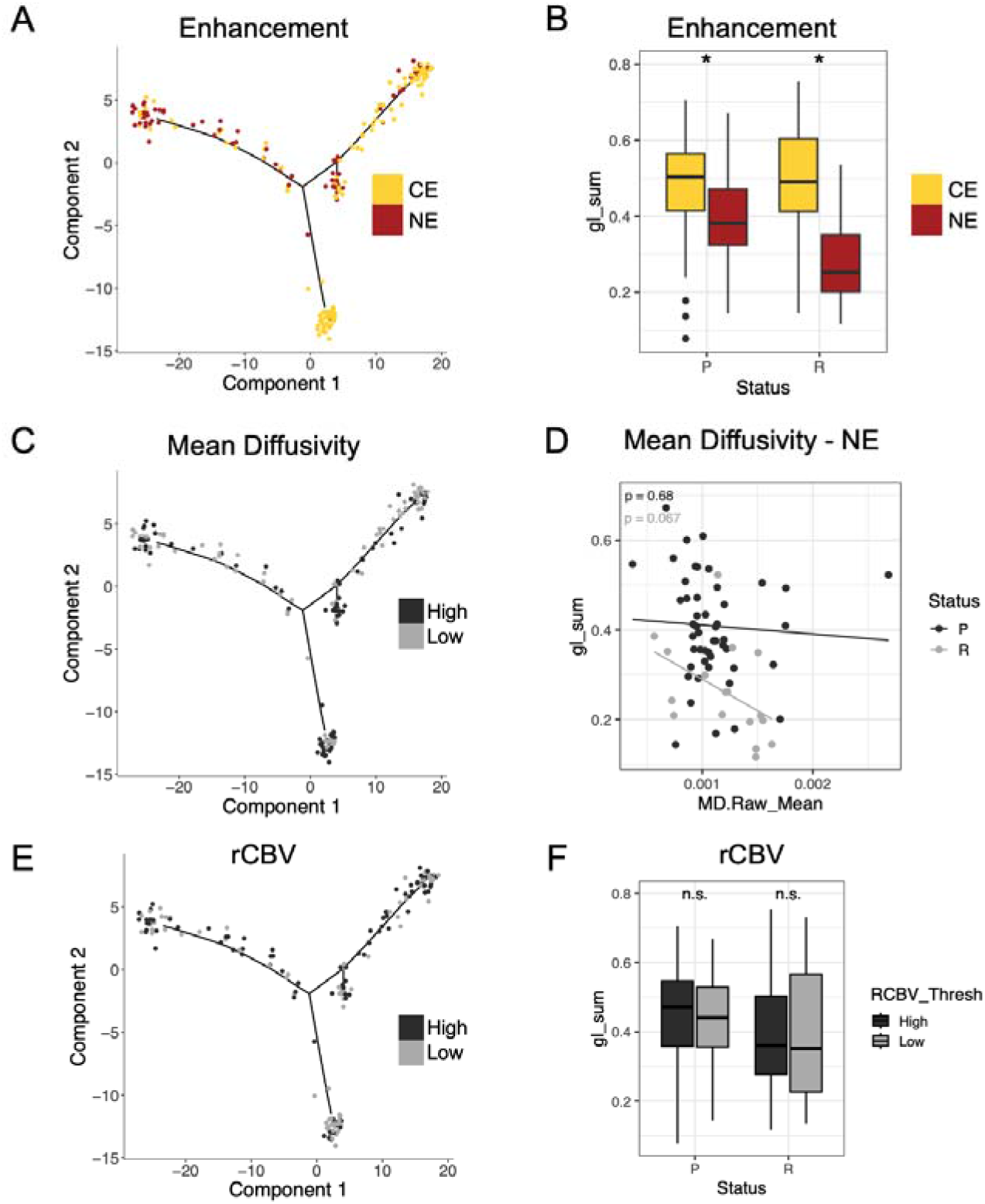
Despite prominent use in the clinic, key clinical imaging features from univariate MR imaging phenotypes fail to distinguish tissue states (arms of the Monocle) and tumor enrichment (gl_sum). **(A)** NE samples dominate the diffusely invaded brain State 3 (left arm) whereas CE did not differentiate immune/inflammatory State 1 (top arm) from proliferative/classive State 4 (bottom arm), despite State 4 containing predominantly CE samples. **(B)** CE and NE significantly distinguish gl_sum in both primary and recurrent samples. **(C)** MD did not distinguish Monocle states. **(D)** MD in the NE regions did not significantly correlate with glioma abundance, despite showing a clearer signal in the recurrent than the primary setting. **(E)** rCBV also did not discriminate between glioma states. **(F)** Although used clinically to detect regions potentially enriched in tumor, rCBV also did not discriminate against samples with high glioma abundance, gl_sum, in this dataset.

### rCBV fails to discriminate regions of glioma enrichment

Perfusion MRI is used clinically to differentiate potential regions of regional cerebral blood volume, rCBV, which is used clinically to help identify regions potentially enriched in glioma cells at recurrence, particularly in regions of NE. In our cohort, rCBV did not differentiate glioma cell fraction in the biopsy sample in neither the primary nor the recurrent setting (Figure 3F).

### MRI habitats reveal distinct patterns of cell co-habitation and tissue state

When parsing the individual habitats visible on multiparametric MRI (CE/NE and high or low on MD, FA, and rCBV), unique patterns of cellular cohabitation and function can be identified (Figure 4). Of the 16 possible combinations, 14 were seen with more than 1 sample in our cohort Figure 4 orders the resultant imaging habitats according to the average pseudotime for samples within that habitat. A mean intraoperative registration error of 1.29mm was observed across 21 surgeries (60 samples) with available information (Figure S13A) and no significant differences were observed in MD, FA or rCBV values at biopsy locations between collection sites (Figure S13B). Within a leave-one-patient-out analysis, sample classification within habitats remained unchanged in 7881/8016 iterations (98.3%), with only 19 samples changing habitats (Figure S14A-B). In this analysis, pseudotime habitat ordering showed a robust mean absolute rank displacement of 0.06-0.61 across the habitats (Figure S14C). The lowest habitat numbers are associated with the lowest pseudotime and thus are enriched in normal brain cell populations like Ast1, Neurons, and proneural glioma cells (gl_PN1 and gl_PN2). Habitat 14 is such that all glioma subpopulations are represented, there are minimal-to-no OPCs, neurons, or oligodendrocytes but the region is still enriched in T cells compared to most other CE regions. Habitat 8 is an intriguing perivascular immune niche associated with an abundance of vascular-associated prTAMs which is coordinated with an enrichment in endothelial cells and mesenchymal glioma cells (gl_Mes2). Thus, each imaging habitat is associated with somewhat distinct patterns of glioma biology.

**Figure 4.**
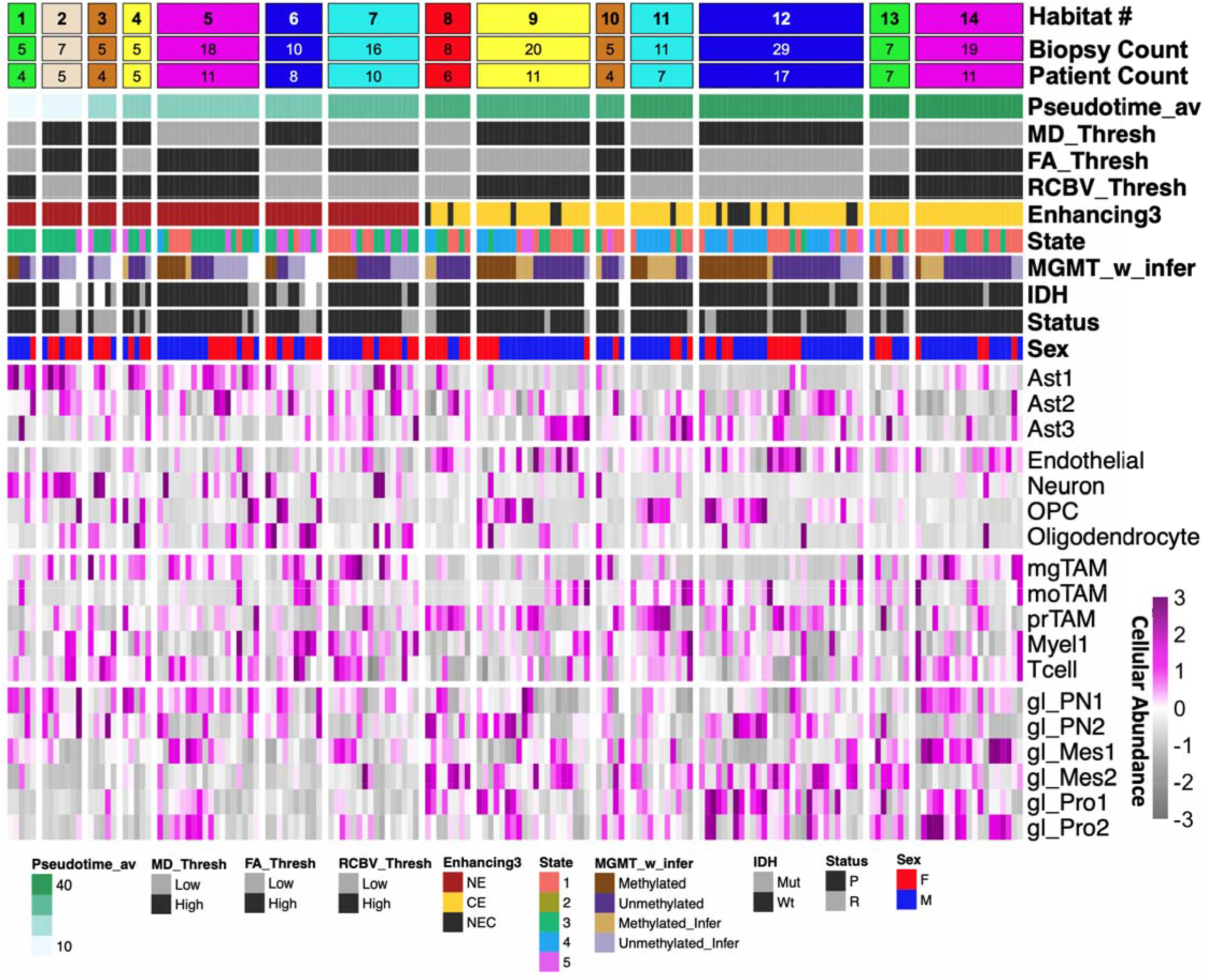
Multiparametric MRI habitats resolve distinct underlying tumor biology. Heatmap of patterns of cell subpopulation cohabitation as a function of imaging habitats defined by multiparametric MRI. Habitats are ordered by the average Monocle pseudotime of biopsies samples associated with each region. This allows direct mapping of the MRI habitat along the continuum of HGG biology/evolution in Figure 2.

### Overlay of MRI habitats on exemplar responder and non-responder patients revealed unique patterns

We mapped these multiparametric MR imaging habitats onto the MRI for our exemplar patient in Figure 2H (Patient 33) in Figures 5A and 5B as well as habitat overlays for an exemplar MGMT unmethylated patient in Figure 4C (Patient 41). Patient 33 was a 59-year-old female with IDH1wt and MGMT-methylated disease who underwent gross total resection followed by standard-of-care radiochemotherapy and had no imageable disease at 4-month follow-up, whereas Patient 41 was a 66-year-old male with IDH1wt and MGMT-unmethylated disease who underwent the same treatment but developed a new peripheral lesion consistent with early progression at approximately 4 months. Habitats 6 and 9 dominated the NE and CE components of Patient 33. Habitat 6 in the NE was notable, amongst MGMT methylated patients like our exemplar patient, for very few tumor cells (low gl_sum) and high abundance of normal Ast1, neurons, and oligodendrocytes. On the other hand, the dominant Habitat 9 in the CE was associated with an enrichment in OPCs, gl_PN2, gl_Pro1 (for MGMT methylated samples). Notably, the key MRI differentiator separating NE dominant Habitat 6 and CE dominant Habitat 9 was rCBV switching from low to high, which was perhaps consistent with the near significant difference in vascular associated signal in terms of endothelial cells (p=0.058). That said, for patient Patient 41, Habitat 6 is similarly prominent in the NE whilst there is a significant increase in the relative abundance of Habitats 12, 13, and 14 in the CE (arrow - Figure 4C). These habitats had more gl_PN1, gl_Mes1, and gl_Pro2 than Habitat 9 which dominated the CE of Patient 33. Habitats 12, 13 represented an admixture of both tissue states 1 and state 4 whilst the increased abundance of Habitat 14 was more suggestive of immune/inflammatory tissue state 1.

**Figure 5.**
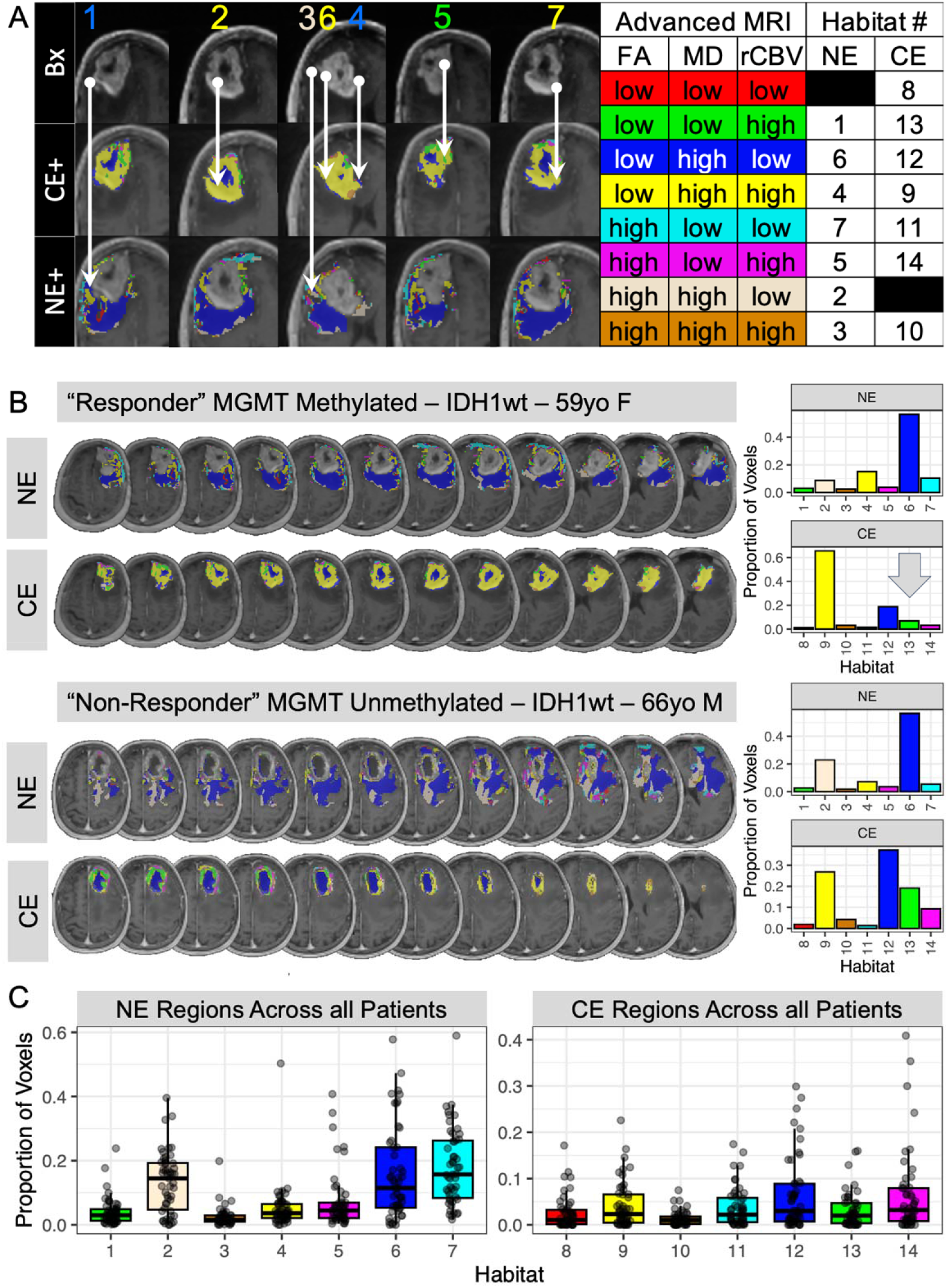
Expanding MRI habitats across abnormal regions. (A) Habitat regions overlaid onto the T1Gd MRI of the exemplar patient from Figure 2H - Patient 33. Top row of the left subpanel shows MRI slices with known biopsy locations, with the middle row showing the CE+ habitat overlay, and bottom row the corresponding NE+ habitat overlay. A range of habitats are seen to be present, with Habitat 6 (low FA, high MD, low rCBV) dominating in NE+ and Habitat 9 in CE (low FA, high MD, high rCBV). (B) Comparison of (top) responder Patient 33 59yo F GBM with MGMT methylation (no imaging progression after SOC) and (bottom) similar located and sized lesion from a non-responder Patient 41 (66yo M with unmethylated MGMT) with significant imaging progression immediately after SOC. Multislice overlay of the MR habitats on methylated exemplar patient (top) and unmethylated patient (bottom) showing an increase in habitats associa ed with higher pseudotime (12-14 - grey arrow) and an associated more inflammatory/immune tissue state. (C) Proportion of voxels found across the cohort of 49 patients with advanced imaging available.

### Low pseudotime MRI habitats tend towards the leading edge of the imaging abnormality

Low pseudotime habitat numbers (associated with low abundance of glioma cells) were generally found at the periphery of the imageable tumor in the NE regions. That said, NE MRI habitats reflected variable states of tumor biology and cellular composition. For example, Habitat 1 through 4 were associated with the lowest pseudotime diffusely invaded glioma state 3. Samples in these habitats had relatively low tumor abundance (gl_sum low) and these habitats were found in patches at the far periphery “invading edge” of the imaging abnormality (Figure 5A). These habitats were coordinately enriched in normal brain cell subpopulations (Ast1, Neurons) and proneural glioma cells (gl_PN1). Some habitats were similarly located at the far periphery of the NE imaging abnormality (e.g. Habitat 5 in Figure 5A bottom panel) that was variably enriched in tumor cells (compared to say Habitat 1), and had other tissue states represented, including the highly reactive/inflammatory State 1. Compared to Habitats 1-3, Habitats 4 and 6 were still NE regions, but they were more closely proximal to the margin of the CE region and had associated higher pseudotimes reflecting a progression along a biological trajectory from tumor periphery to core towards either the more proliferative State 4 (bottom arm) or the more immune/inflammatory State 1 (top arm).

### MRI habitats differentiated the glioma ecosystem into biologically meaningful ecologies

Jensen-Shannon divergence (JSD) quantifies the degree to which imaging regions were divergent from each other in terms of cellular composition (Figure S12). The most discriminatory univariate MRI region was CE vs NE with a JSD of 0.032, compared to 0.008 for MD high vs low, 0.014 for FA high vs low, and 0.005 for rCBV high vs low.

### MGMT methylation status coordinates with differences in patterns of cellular co-habitation within habitats

Figure 4 illustrates a notable distinction within each habitat related to MGMT methylation status. Consistent with this, unmethylated samples were in the higher pseudotime immune/inflammatory glioma state 1 (top arm compared to the more proliferative state 4 - bottom arm - Figure S4B) for CE samples, there was comparable shift in higher pseudotime associated MRI habitats (12, 13, 14) for the unmethylated patient (bottom) than the methylated (top) - arrow in Figure 5B. Notably, in some of the most well sampled habitats, MGMT methylation is associated with increased OPC abundance, like in habitats 10, 11, and 12 (p=0.035). There is also an increase in gl_Pro1 and gl_PN2 in MGMT methylated samples in habitat 9 (p=0.00053, p<0.0001, respectively).

### Prognostic Significance of MRI Habitats Associated with Aggressive Tumor Biology

When applying this MRI habitat method across the patient cohort with advanced imaging and available overall survival (49 patients, Figure 5C), univariate Cox proportional hazard models revealed two significant habitats by voxel counts, Habitat 9 (p=0.039) and Habitat 11 (p=0.009) (Figure 15A). Habitat 10 showed an insignificant but notable signal (p<0.1). As expected, age at diagnosis and IDH status were both also prognostic (p<0.001, p=0.03, respectively), however MGMT status alone was not (p=0.44) (Figure S15B). Multivariate analyses combining all three of these habitats were implemented with age at diagnosis and additionally on the subset of patients with available MGMT, and IDH status. Habitat 11 remained significantly prognostic with both age at diagnosis (p=0.029, Figure S15C) and age at diagnosis with IDH status (p=0.03, Figure S15D). Habitat 9 was also significant when adjusting for age at diagnosis and MGMT (p=0.038) (Figure S15E). FGSEA analyses comparing Habitat 9 and Habitat 11 to all others show that they are both upregulated in different cancer-related pathways. Habitat 9 is associated with upregulated proliferation pathways and Habitat 11 showing immune response, and signaling pathways, amongst others (Figure S16).

## Discussion

Clinical neuro-oncology is challenged by access to brain tumor tissue thus limiting our day-to-day understanding of the heterogeneous tumor spatial and temporal landscape. Much of our insights from clinical imaging have been driven by scant studies matching usually a single patient sample to a single patient MRI with no image localization. To address the lack of MRI-to-biology mapping in glioma research, we leveraged a novel cohort of multiregional image-localized biopsies to connect locoregional MRI features with transcriptional states and cellular composition across 58 patients. Our work can begin to overcome the common limitations of solitary non-localized sampling.

To simplify the profound complexity across samples, an unsupervised trajectory inference algorithm revealed that the HGG transcriptome from our multiregional samples (Figure 2G) and TCGA (Figure S5) could be ordered with a pseudotime approach along a continuum connecting three polarized glioma tissue states: invasive (left arm - State 3), immune/inflammatory (top arm - State 1), and proliferative (bottom arm - State 4). The trajectory’s constrained nature may have implications for treatment, as it proposes the possibility that transcriptional dynamics could be reversed or shifted along the graph toward a more favorable (less aggressive or treatment amenable) state. This enhances the potential for adaptive therapy schemas^44^ which are increasingly focused away from cytotoxic treatments and towards cytostatic population control approaches. Figure 2G provides a visually compelling illustration that transitions between glioma tissue states reflects a coordination of shifts in cellular subpopulations. Low pseudotime is dominated by normal brain cells (protoplasmic astrocytes, neurons, and oligodendrocytes) towards moderate pseudotimes, which are associated with the bottom arm of the Monocle glioma tissue state 4, showing a coordinating influx of OPCs and proneural (gl_PN2) as well as proliferative (gl_Pro1 and gl_Pro2) subpopulations with some coordination of vascular-remodeling associated prTAMs. Then, as pseudotime progresses along the increasingly mesenchymal inflamed/immune glioma state 1, we see an influx of immune cells (Tcells, monocyte-like moTAMs, myeloid cells - Myel1, reactive astrocytes - Ast3, vascular endothelial cells and mesenchymal tumor cells (gl_Mes1 and gl_Mes2). These states also showed some prognostic promise when characterizing TCGA microarray data, particularly when comparing patients characterized as State 1 against those as State 4, a result that was significant amongst female patients. For now, we consider this result as hypothesis generating, as this dataset only has one sample per patient and overlooks the vast intra-tumor heterogeneity we have demonstrated. We aim to verify these results in a more appropriate larger image-localized cohort.

Others have sought to map tumor biology to these overall imaging metrics including the original study in 1987^45^ as well as more recent deeper dives^8^. In our study, samples from regions of CE were further along the biological progression trajectory with higher pseudotime than samples from NE regions. This supports the premise that NE regions will biologically progress to become regions of CE on future imaging follow-up, which is certainly consistent with patterns of serial imaging follow-up of patients^46,47^. Further, CE was associated with increasing abundance of malignant cell populations and was associated with higher expression of many cancer hallmarks. These findings in isolation support the traditional notion that CE and NE represent proliferative tumor and invaded brain, respectively. However, in our data CE alone had little ability to discriminate between the more immune/inflammatory tissue state from proliferative glioma state (Figure 3A). Yet, distinguishing these tissue states is of utmost importance in distinguishing not only understanding treatment response in patients but also for tailoring treatment that best targets said biological state. For example, cytotoxic strategies will likely have more impact on the bottom arm proliferative glioma state 4 whilst immunotherapies would have potential T-cells to ignite in top arm state 1. A similar analysis of diffusion and perfusion MRIs alone also failed to distinguish clinically relevant features of HGG biology (Figure 3). It was particularly unexpected to see that rCBV did not significantly distinguish glioma cell fraction (gl_sum) in the recurrent setting (Figure 3F), yet this may be due to the relatively small sample size of this subcohort (38 samples, 12 patients) collected across two institutions

That said, by combining multiparametric MRIs into regional habitats, we revealed distinct patterns of these habitats across each patient’s brain (Figure 4). By ordering the MRI habitats by the average pseudotime of samples from such habitats, we found that low pseudotime, diffusely invaded brain samples logically appear near the periphery and increase in pseudotime toward the center of the lesion. We also observe some prognostic impact of these habitats (Figure S15), with higher levels of Habitat 9 and Habitat 11 both showing significantly worse overall survival outcomes in some settings and associations with aggressive cancer hallmark pathways. However, these prognostic findings need further validation on larger image-localized biopsy cohorts. Both the sample classification within habitats and the habitat pseudotime ordering were robust to leave-one-patient-out perturbation. This builds the framework for connecting regional MRI habitat distributions to an overall schematic of tumor progression from the leading-edge periphery towards the central core of the lesion.

Immunotherapy has shown strong promise and results as a treatment across a plethora of different cancer types. Historically, the brain was thought to be separated from the immune system, but it has since become clear that the immune system is indeed present and interacting within the brain^48^. In a similar vein, glioma has been considered “cold” with little lymphocytic infiltration, but T-cells and myeloid cells (including tumor-associated macrophages) are present and can account for up to half of a sample^49^. It is thus of great interest whether similar success for immunotherapy can be seen in glioma, although to date no paradigm-shifting clinical trial results have emerged^50^. We were interested in the spatial localization of these immune populations, how they cohabitate with glioma subtypes and other cell types. Notably, only certain MRI habitats showed enrichment for immune populations that may be prime targets for reprogramming of the immunosuppressive “cold” environment into a “hot” immunoresponsive environment. For instance, Habitat 14 showed not only an enrichment in T-cells, but also a coordinated expansion of other immunosuppressive TAM and myeloid populations. These MRI habitats may be highly relevant in future studies of response in immuno-oncology trials.

We presented a case example of a patient who underwent seven biopsies from both CE and NE regions (Figure 2H, Figure 5A). Despite cohort-wide trends of cellular composition and contrast enhancement, the cellular composition of these samples was not predicted by contrast enhancement status as we would have assumed. This result highlights the limitations of current methods of treatment assessment, where a tumor is considered to be responsive if the CE volume regresses. The inability of contrast enhancement alone to fully predict cellular composition motivates a role for advanced imaging strategies like radiomics. Radiomics is a quantitative approach to predict biological attributes from complex imaging features ^51,52^. Early radiomics efforts focused on connecting an entire image to a biological prediction, thus failing to capture the spatial heterogeneity within individual tumors. More recently, our group and others have leveraged image-localized biopsies to directly connect spatially-resolved imaging features with tissue biology ^28,29,53–56^. Sex differences in transcriptional expression and its relation to bioimaging embolden the careful incorporation of these variables in the context of radiomics. Elucidating these connections will provide the opportunity to non-invasively characterize tumors in their entirety and personalize each patient’s therapy to exploit the unique weaknesses of their tumor’s dominant state.

Lastly, by overlaying the multiparametric MRIs onto each other and breaking down MRI regions into habitats of high or low intensity across multiparametric MR sequences, we were able to identify 14 unique imaging habitats represented in our MRI-localized data set. Numerous prior studies have considered imaging habitats as a concept,^57^ but few have had imaging-localized tissue to provide validation of the biology predicted to underlie each imaging habitat^58^ or a few samples across a limited cohort of patients (e.g. 17 patients with multiparametric MRI and PET with 31 biopsies^59^). This is not surprising given the resource-intensive needs of collecting MRI-localized samples clinically and the shifts in clinical workflow needed to collect multiregional samples reliably^59,60^.

Using Jensen–Shannon divergence (JSD)^61–63^, we found that our MRI habitats retained distinct patterns of cellular composition forming biologically distinct ecological niches (Figure S12). Some of the MRI habitats had more distinctive and definable cellular compositions but others may best be characterized by non-compositional characteristics. We found that JSD patterns differed between CE and NE habitats, with greater compositional similarity among samples within habitats than between. A subset of NE and CE habitats exhibited similar compositions across both states, forming a “bridge” between these biological states (Figure S12A). There is a rearrangement of the grouping/similarity of the MRI habitats when considering the JSD for glioma cells subpopulations only, or immune only, or other non-glioma/non-immune (Figure S12B-D). This suggested that composition shifts in these subcomponents could be connected to distinct shifts in MRI habitats patterns in each patient. Comparing JSD within and between MRI habitats showed that there is significantly greater compositional variation between MRI habitats than within (Figure S12E). For these reasons, we were hesitant to collapse habitats together. Additionally, habitats with fewer samples in this cohort may be better represented in others. In a leave-one-patient-out analysis, we observed minimal movement of habitat assignment and ordering (Figure S14). Taken together, these data suggest that MRI-defined habitats exhibit significantly lower within-habitat ecological divergence compared to between-habitat divergence, supporting biologically coherent microenvironmental structuring. These data support a future in which such quantitative imaging approaches could be used to detect biologically meaningful shifts in the tumor ecosystem within each patient.

MGMT methylation is associated with reduced DNA repair capacity, and thus more DNA damage/mutations accrue with treatment^64^. MGMT methylation also tags a tumor that lives in a progenitor-like epigenetic state, often overlapping with OPC-like biology. This is consistent with our observation that MGMT methylation is also associated with remodeling of the tumor microenvironment to enhance enrichment in OPC with habitats. Consistent with prior reports of MGMT unmethylated samples showing increased immunosuppressive markers, MGMT methylated samples appear more prominently on the proliferative tissue state 4 while MGMT unmethylated are more common on the top arm immune/inflammatory tissue state 1 (Figure S3B). ^65^

Many future works are motivated by these results including 1) ongoing collection of large cohorts of MRI localized biopsies to enhance the MRI-biology mapping, 2) coordinated biological studies to probe how tumor biology and thus MRI habitats might perturb as a function of treatment, and 3) additional validation studies to establish the reproducibility of MRI habitats in detecting meaningful shifts in tumor biology.

## Ethics Statement

This work was carried out with approval from the Mayo Clinic institutional review board (16-002424) and the Cedars-Sinai institutional review board (STUDY00004501).

## Funding

National Cancer Institute (U01CA250481 to N.L.T., L.S.H. and K.R.S., U54CA274504 to P.C. and K.R.S., U54CA274507 to A.R.A.A.). Cross-consortium funding from the National Cancer Institute Cancer Systems Biology Consortium (U54CA274504, U54CA274507, to K.R.S. and A.R.A.A.). American Cancer Society Arizona Discovery Boost Award (to K.R.S.). Institutional support for authors, including from Cedars-Sinai Medical Center and Mayo Clinic.

## Acknowledgments

We are grateful to the patients who graciously shared their data. We would like to thank those who have contributed to elements of this work, particularly current and past members of the image analysis teams and the glioma biopsy protocol teams, including: Barrett Anderies, Jessica Bauer, Spencer Bayless, Hend Bcharach, Regina Becker, Sameer Channer, Brenden Doyle, Lysette Elsner, Lily Esaleh, Ashlyn Gonzales, Crystal Harris, Morgan Hatlestead, Ryan Hess, Sandra Johnston, Julia Lorence, Ashley Napier, Ashley Nespodzany, Cassandra Rickertsen, Sejal Shanbhag, Sarah Van Dijk, and Scott Whitmire.

## Conflicts of Interest

Fuzionaire Theranostics (stock: K.R.S.); Imaging Biometrics (medical advisory board: L.S.H.); the remaining authors have no conflicts of interest to disclose.

## Data Availability

All datasets used in this work are publicly available. Bulk RNAseq counts data can be found at Synapse with SynID syn52256644 (metadata batches - D3-D5) and SynID syn74801419 (metadata batches D6 and D6_low). Code, metadata and batch-corrected counts for this manuscript are available in a public repository here: https://github.com/LeeCurtin/HGG_Monocle_CIBERSORTx. The snRNAseq dataset used for CIBERSORTx can be found here: https://github.com/adithyakan/reconvolving_gbm.

## Authorship

LC: Informatics, data collection, data curation, data analysis, interpretation, statistics, writing

KMB: Conception, informatics data curation, data analysis, interpretation, writing

AHD: Informatics, image processing, data analysis

JCU: Image-localized biopsy collection, image processing, data curation

GD: Image-localized biopsy collection, image processing, clinical data abstraction

CS: Biopsy tissue processing

KWS: Image processing

JML: Image visualizations

PRJ: Writing

CK: Data collection

RSZ: Data collection

DPP: Data collection

BRB: Data collection

KS: Data collection

PN: Data collection

KD: Pathology review

LB: Data collection

MMM: Patient consenting, interpretation

CP: Informatics

OA: snRNAseq collection, analysis

LSH: Imaging collection and processing, interpretation

NLT: Tissue processing

JG: Data interpretation

ARA: Results interpretation, writing

JBR: Results interpretation, writing

PC: Data collection, data analysis, interpretation

KRS: Conception, informatics, data collection, data analysis, statistics, writing

## Notes

### Competing Interest Statement

Fuzionaire Theranostics (stock, KRS); Imaging Biometrics (medical advisory board: LSH); the remaining authors have no conflicts of interest to disclose.

### Summary of Updates

Primary additions are a leave-one-patient out analysis, application of MRI habitats across imaging abnormalities cohort wide, and a subsequent overall survival analysis.

