## Supplementary Figures and Tables for "Mapping multiregional image-localized biopsies to MRI habitats reveals biologically significant glioma tissue states and patterns of cellular subpopulations"

### Figure S1. Breakdown of CIBERSORTx predictions by snRNA downsample and population predicted

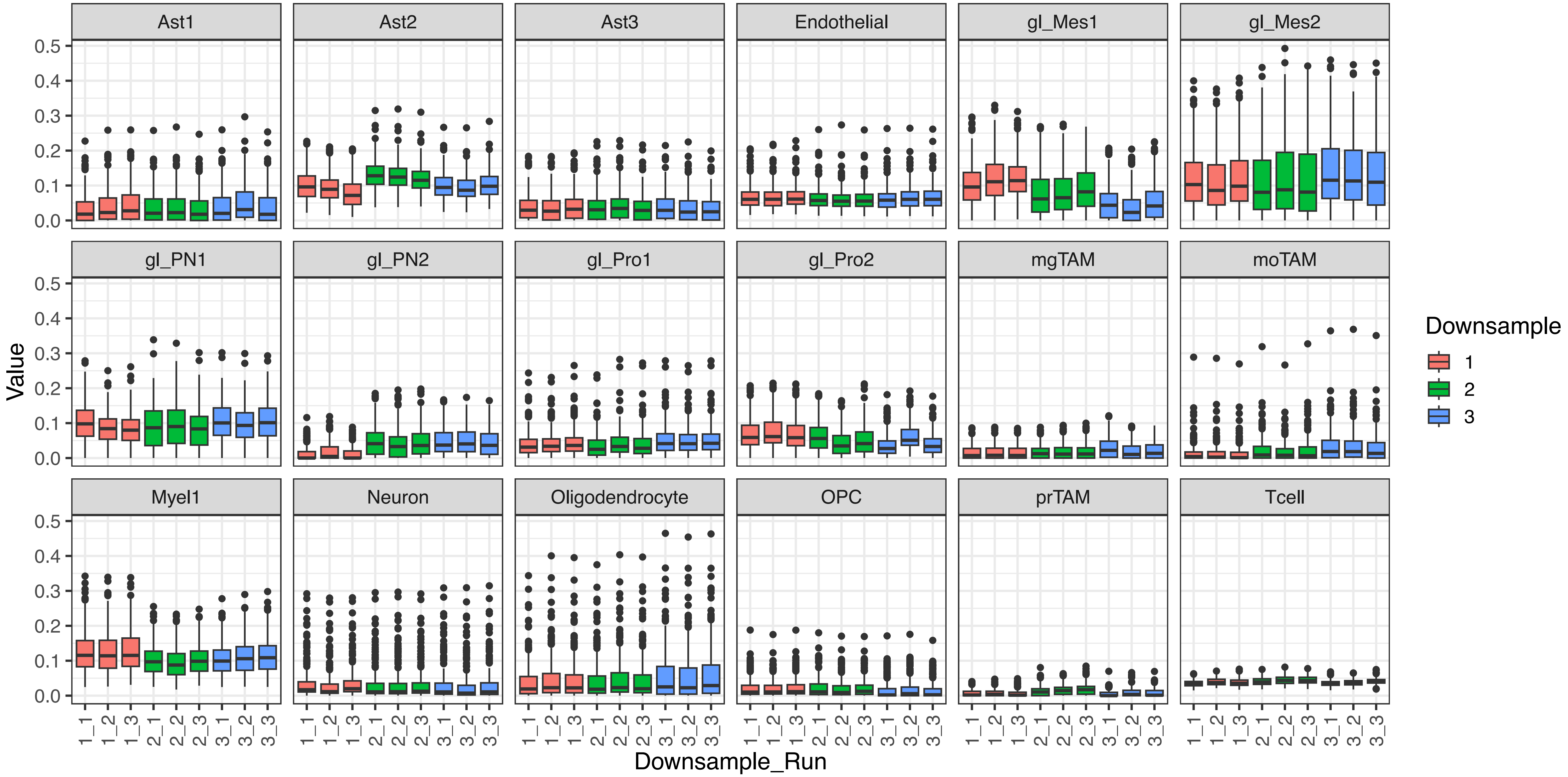

Figure S2: (A) Bulk GSEA comparing each state to all others using cancer hallmarks. (B) Single sample GSEA of cancer hallmarks for each sample

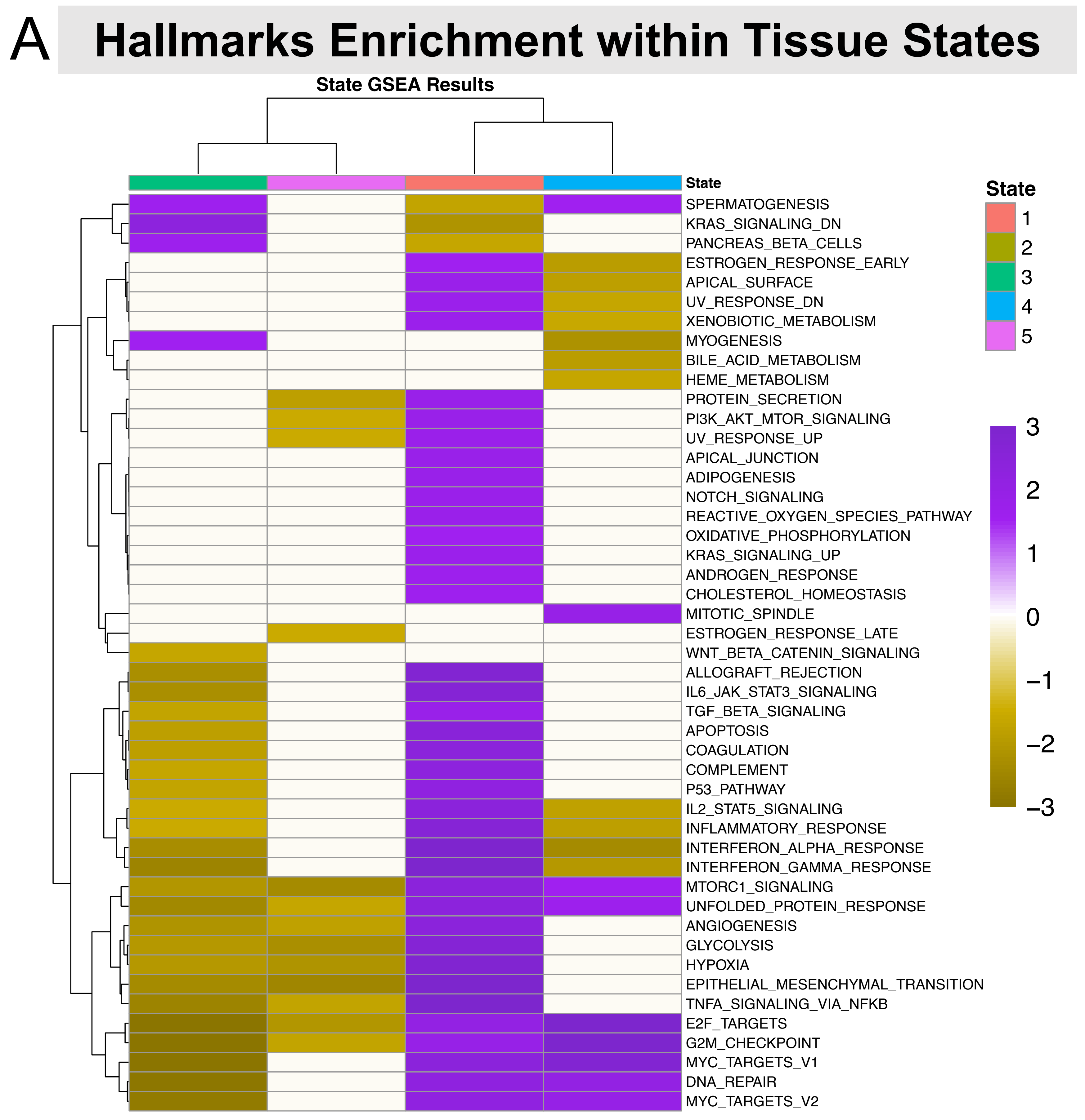


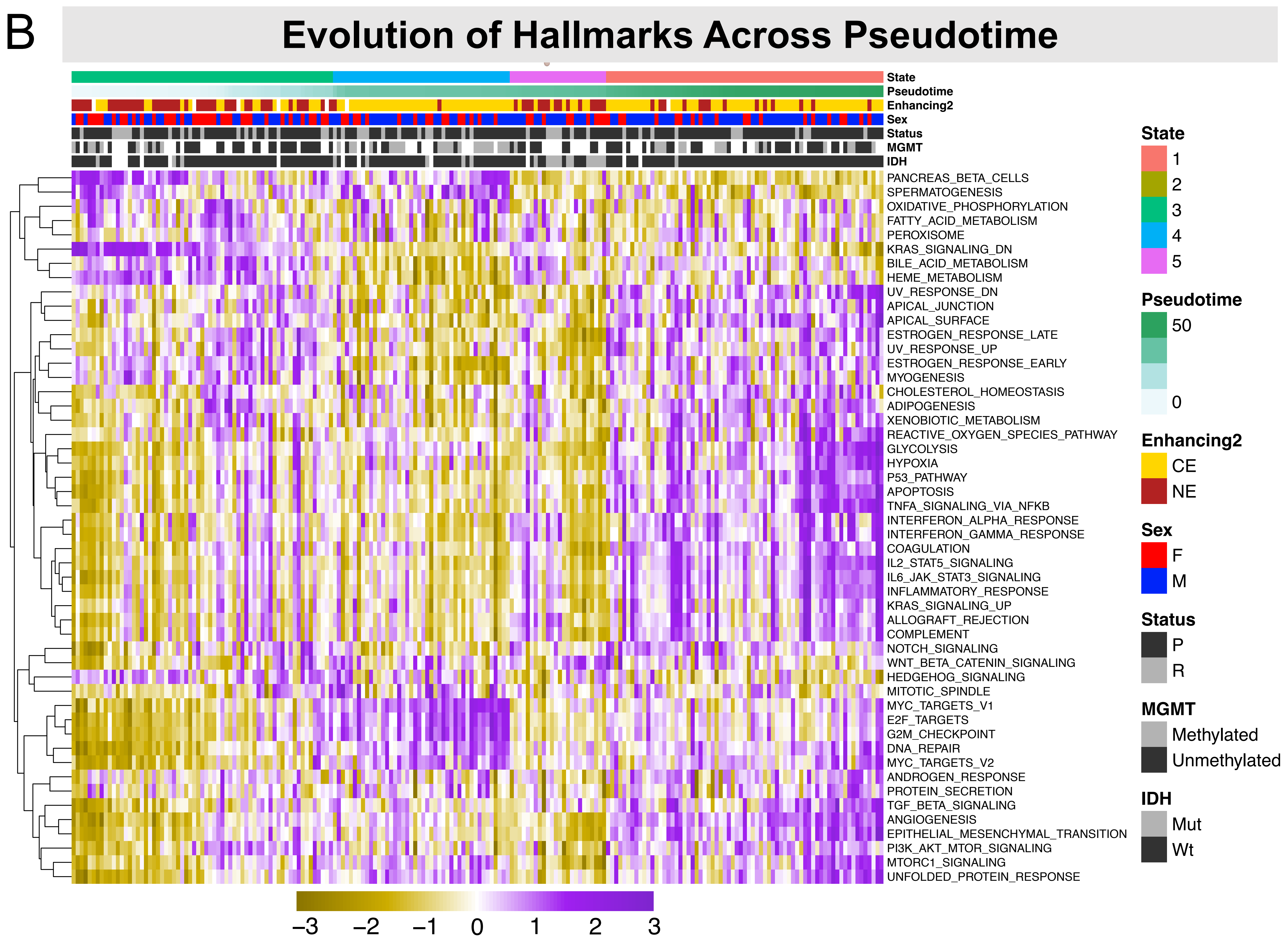

Figure S3: (A) IDH status presented on Monocle graph. (B) MGMT methylation status presented on Monocle Graph

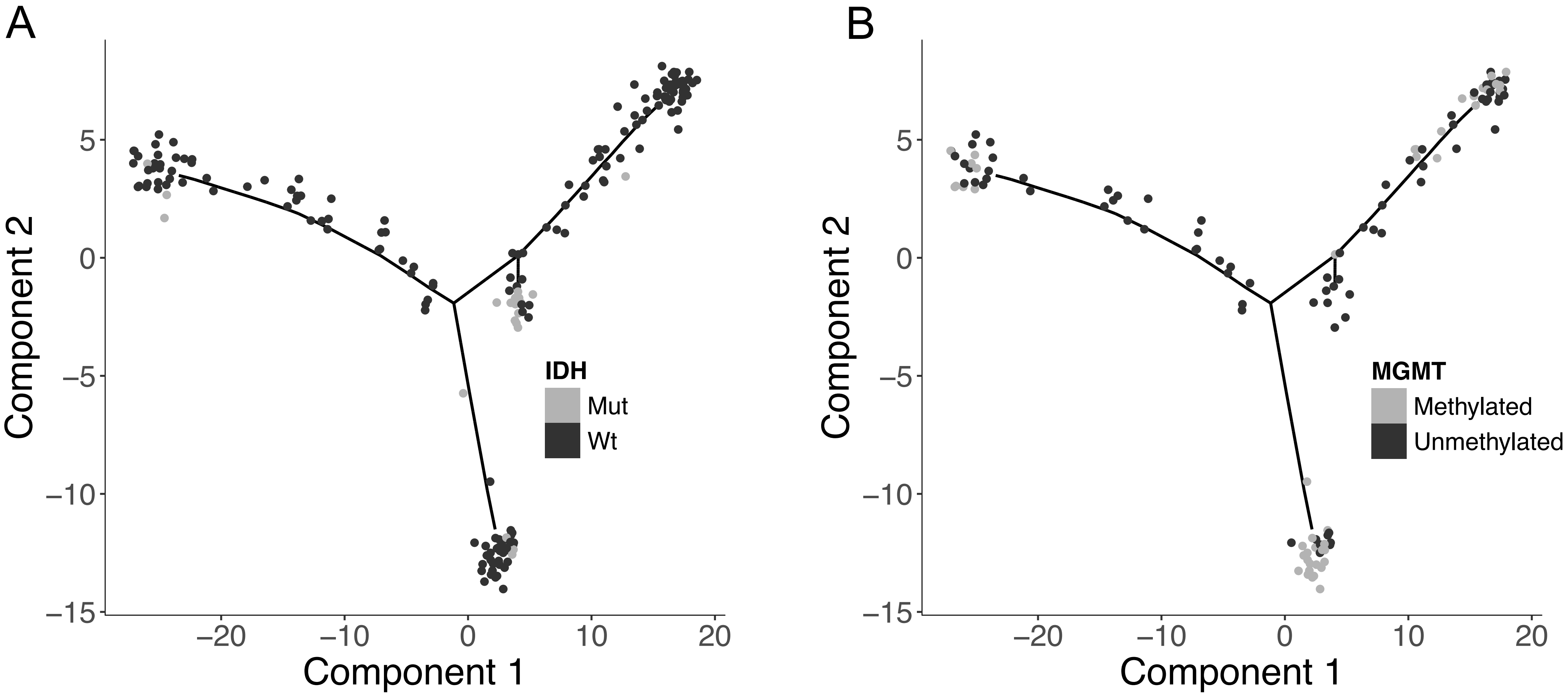

Figure S4: (A) TCGA microarray data shows similar Monocle graph pattern. (B) Monocle graph shows strong associations with pre-classified subtypes

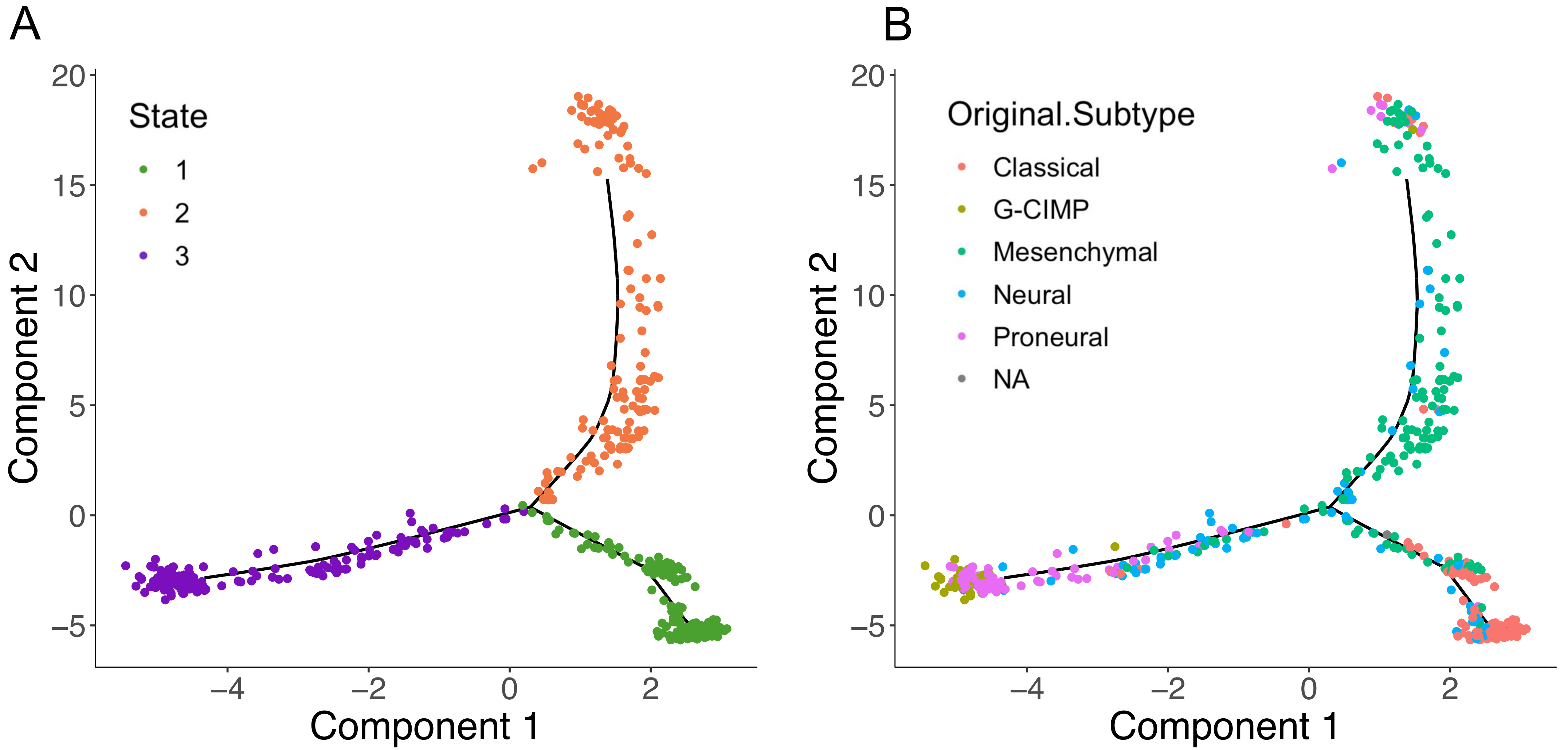

Figure S5: CIBERSORTx-predicted cell states significantly vary with Monocle State

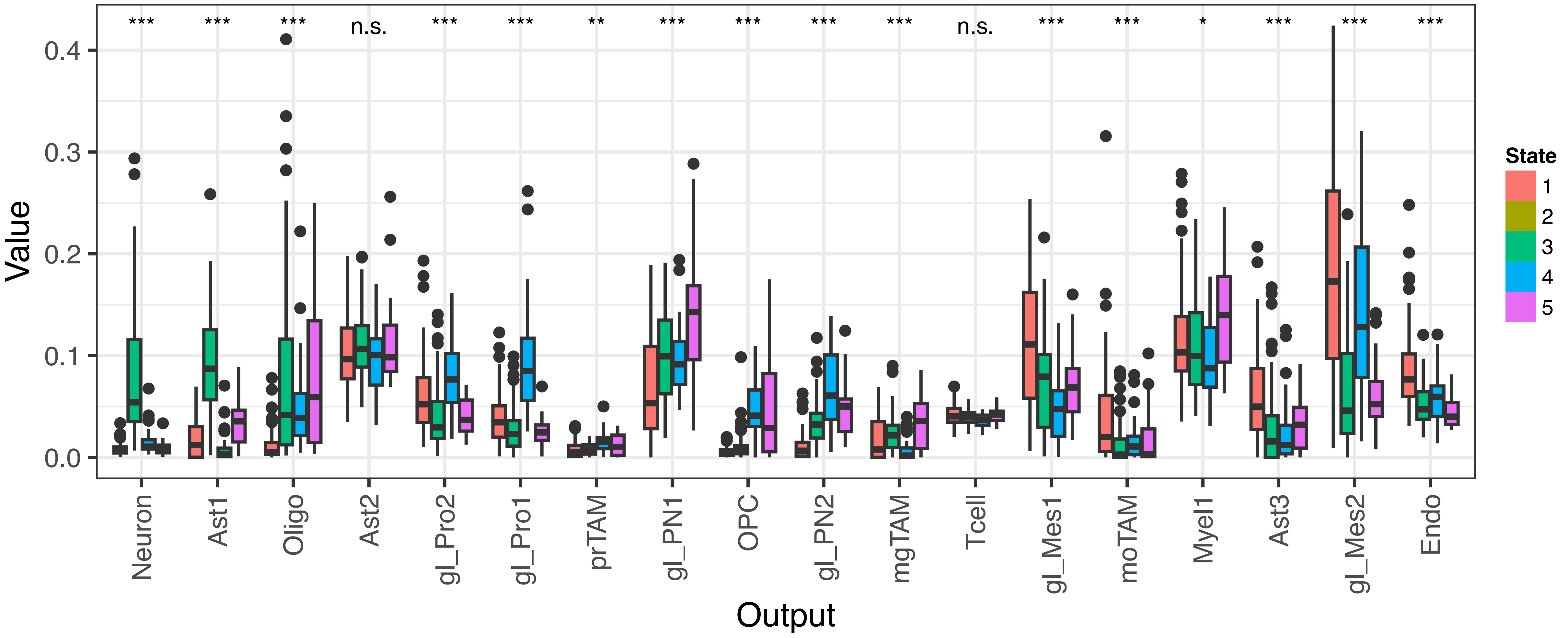

Figure S6: Shannon indices computed across (A) all CIBERSORTx-predicted populations, (B) glioma populations, (C) other populations, and (D) immune populations, presented on an exponential axis for ease of visualization.

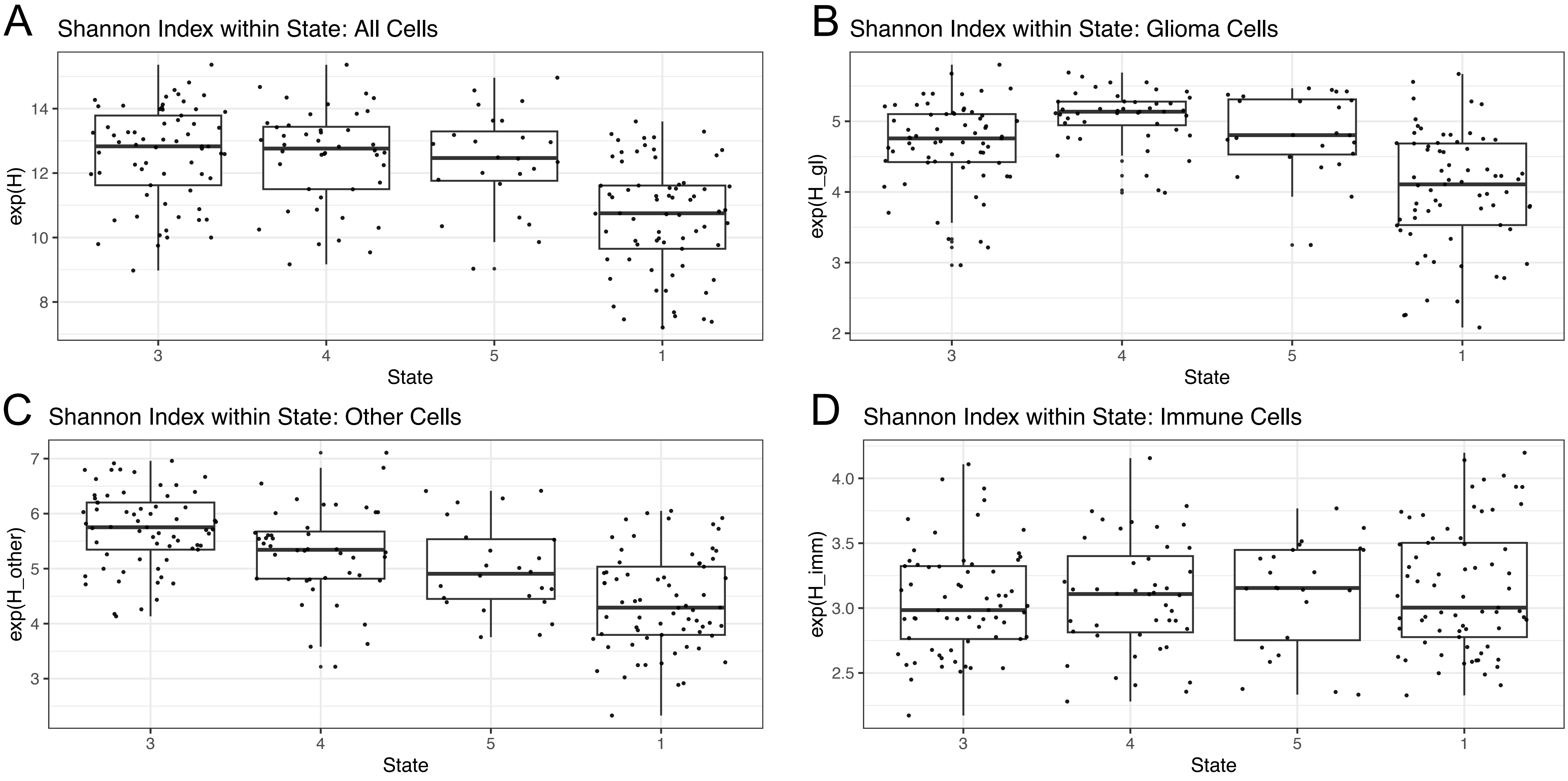

Figure S7: Survival comparing State 1 and State 4 assignments to TCGA microarray data for (A) all samples, (B) female samples (C) male samples.

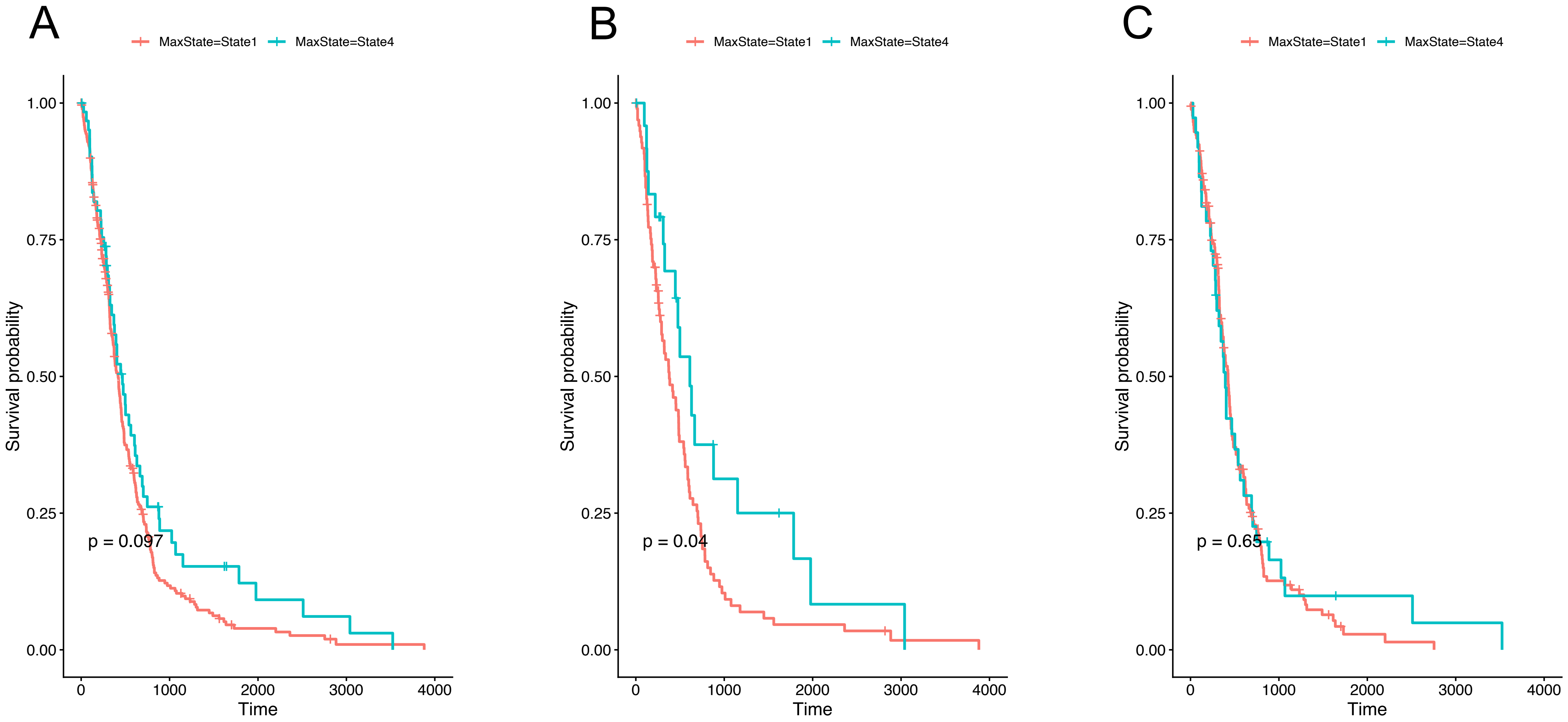

Figure S8: Comparing similarity between samples within Pt33

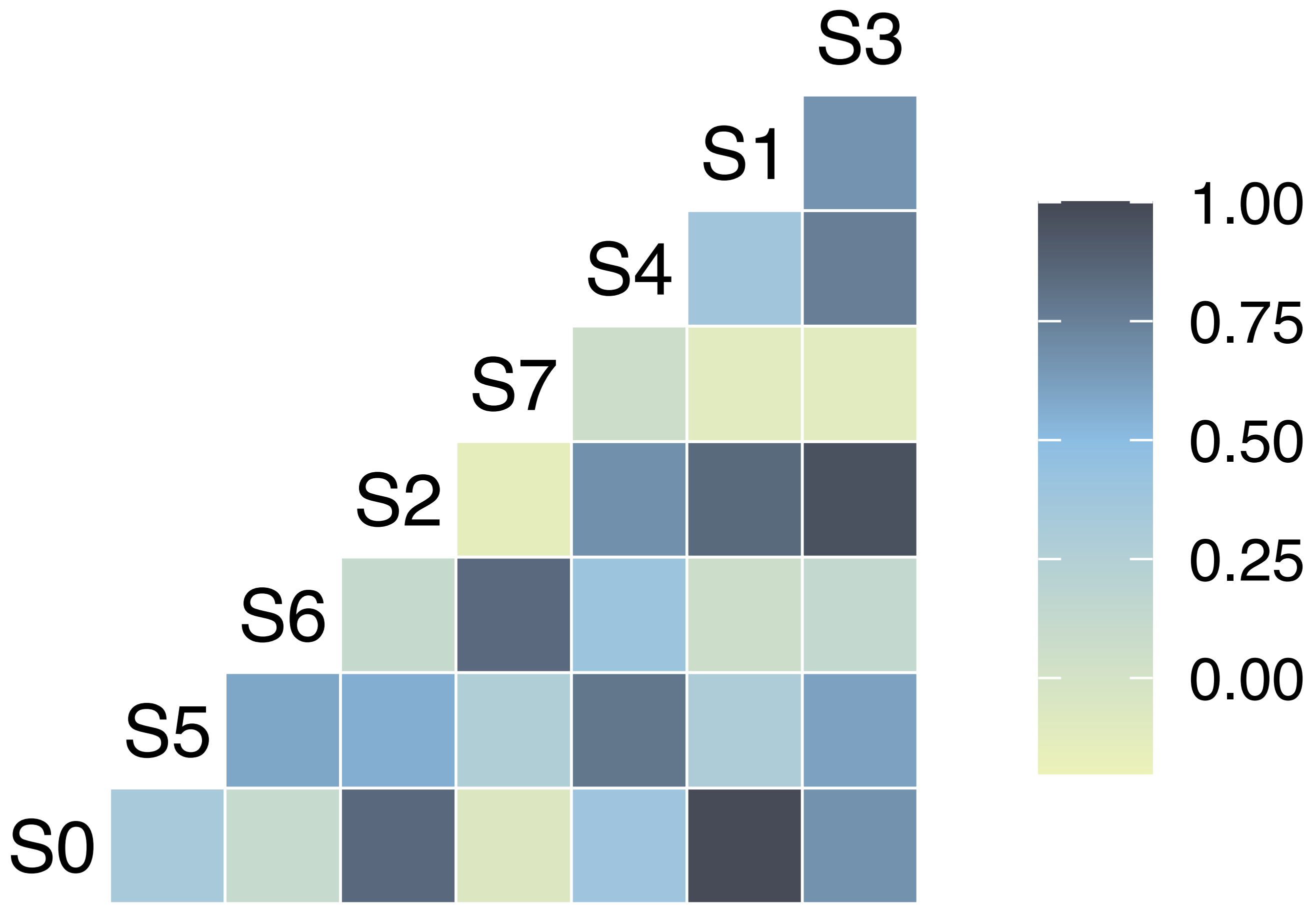

Figure S9. Breakdown of CE into those from the internal edge shows clear grouping towards Monocle end states

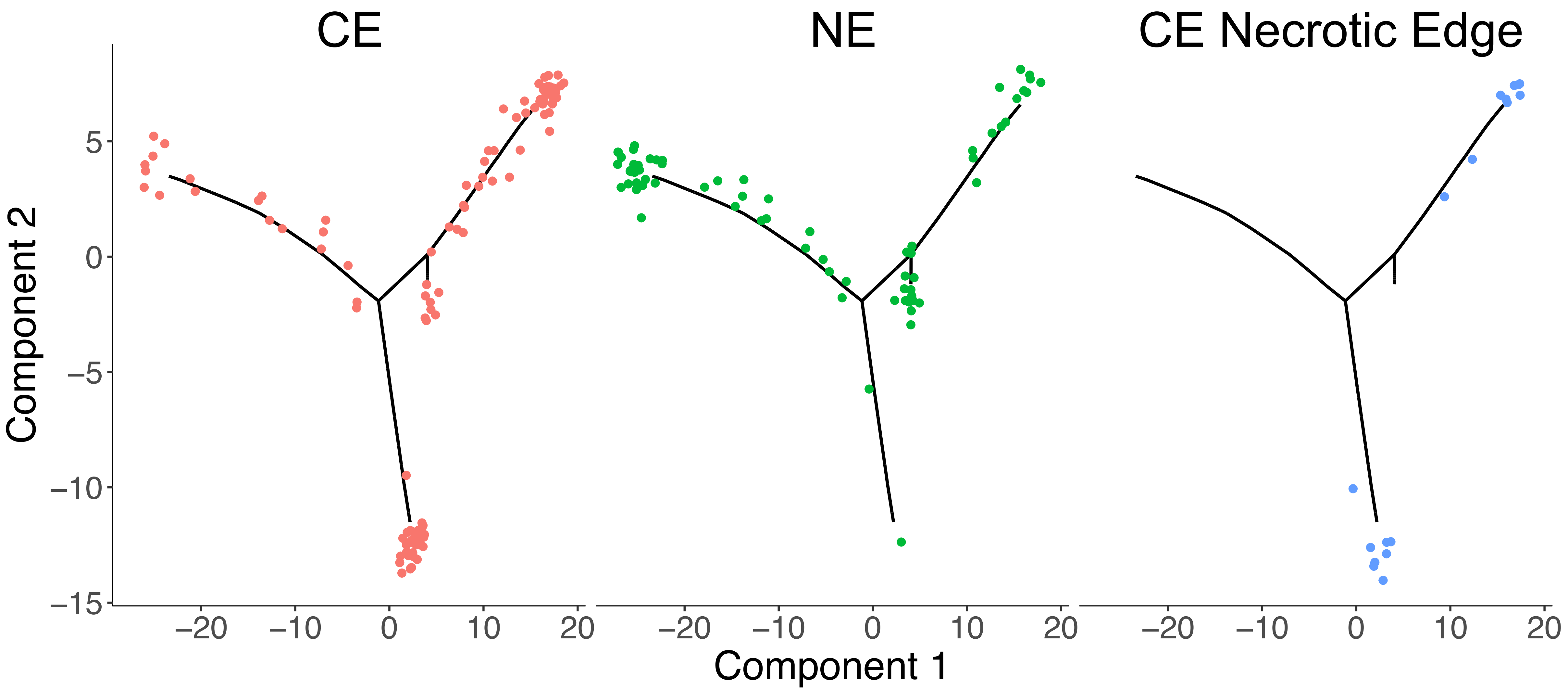

Figure S10. Monocle graph with treatment status overlaid, no clear signal emerges for primary vs recurrent samples

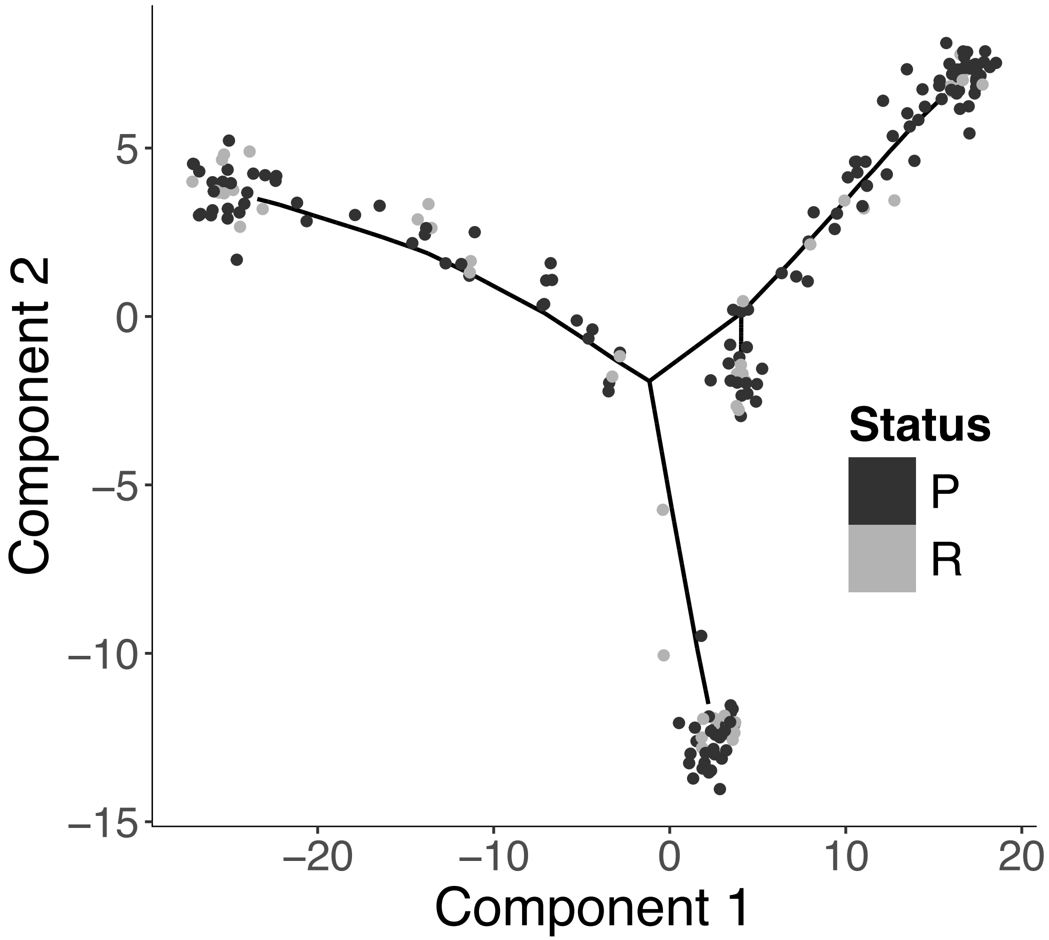

Figure S11: CIBERSORTx-predicted cell states significantly vary with enhancing status

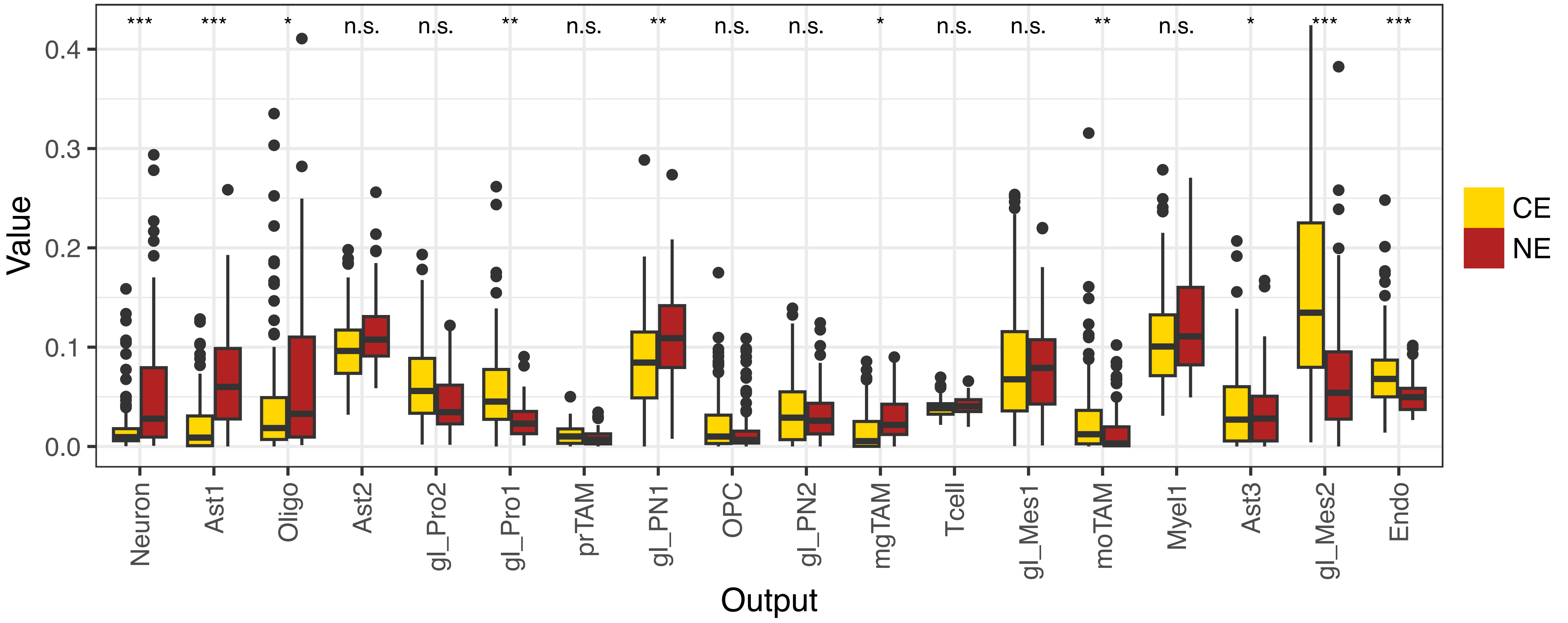

### Shannon- Jensen Diversity Figure S12

4/27/26

**Figure S12.** Jensen-Shannon divergence (JSD) between MRI habitats. A. Divergence amongst all tumor cells shows a clustering of CE and NE habitats as there is more similarity between samples within those habitats rather than between habitats. There appears to be a subset of NE and CE habitats that are similar to habitats in both NE and CE habitats forming a ”bridge” between these biological states B. Divergence amongst glioma cells reveals different associations of MRI habitats in terms of similar glioma cell compositions. Similar for immune cells (C) and other non-tumor, non-immune cells (D). E. Comparing JSD within and between habitats shows that there is more compositional variation between MRI habitats than within. Performing 1000 permutations produces a mean difference between the within and between habitat JSD as 0.021 ($p<0.001$). This supports the hypothesis that MRI habitats are biologically meaningful ecological niches. F. Some habitats are strongly distinguished by their cellular composition while others are less able to be discriminated from other habitats according to cellular composition alone. Habitats 2, 5, 6, 7, 8, 11, 12, 13, 14 are biologically distinct in terms of cellular composition from other Habitats.

S12A

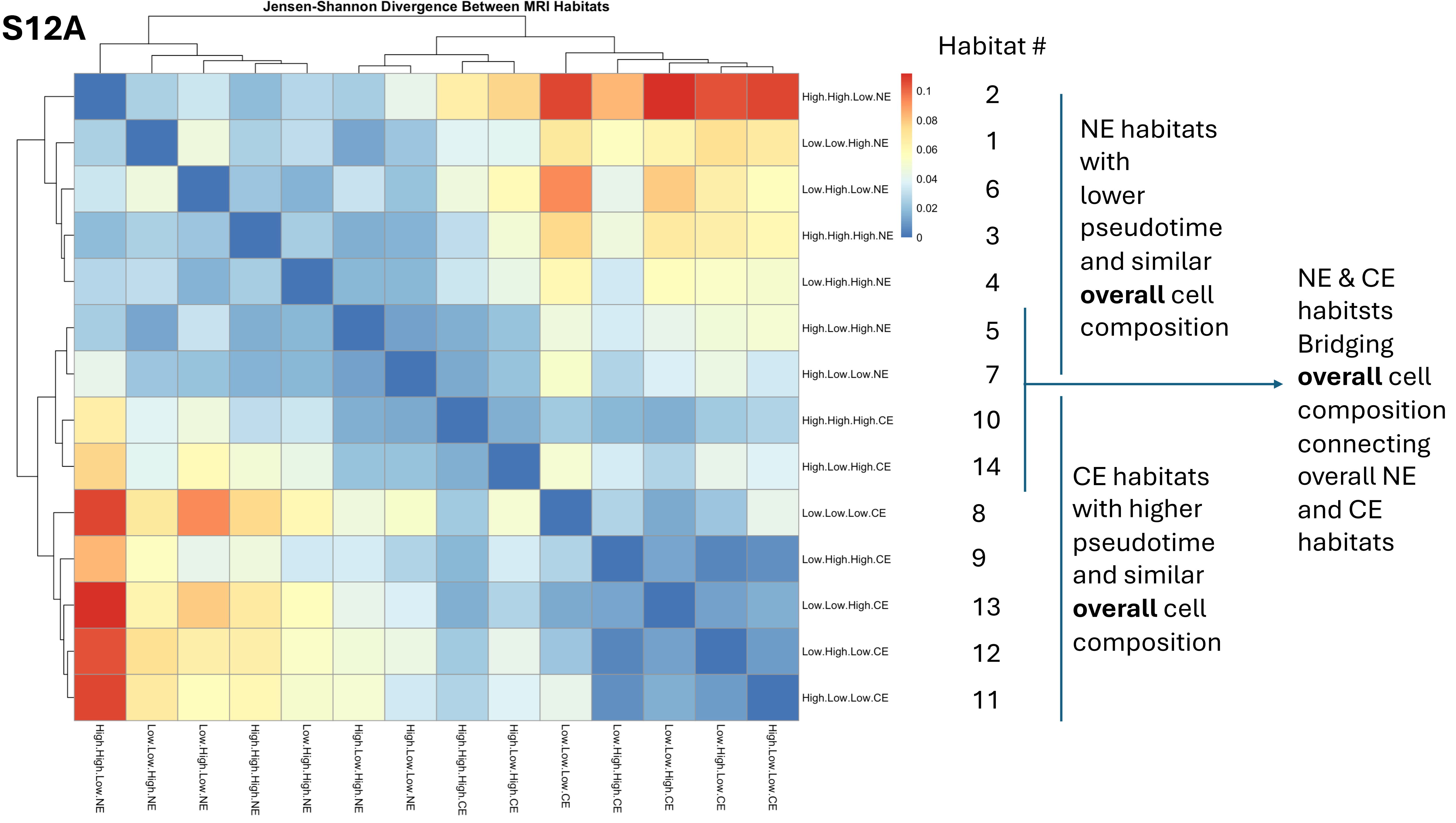
Jensen-Shannon Divergence Between MRI Habitats
High.High.Low.NE
Low.Low.High.NE
Low.High.Low.NE
High.High.High.NE
Low.High.High.NE
High.Low.High.NE
High.Low.Low.NE
High.High.High.CE
High.Low.High.CE
Low.Low.Low.CE
Low.High.High.CE
Low.Low.High.CE
Low.High.Low.CE
High.Low.Low.CE
0.1
0.08
0.06
0.04
0.02
0
Habitat #
2
1
6
3
4
5
7
10
14
8
9
13
12
11
NE habitats with lower pseudotime and similar **overall** cell composition
CE habitats with higher pseudotime and similar **overall** cell composition
NE & CE habitsts Bridging **overall** cell composition connecting overall NE and CE habitats

**S12B**

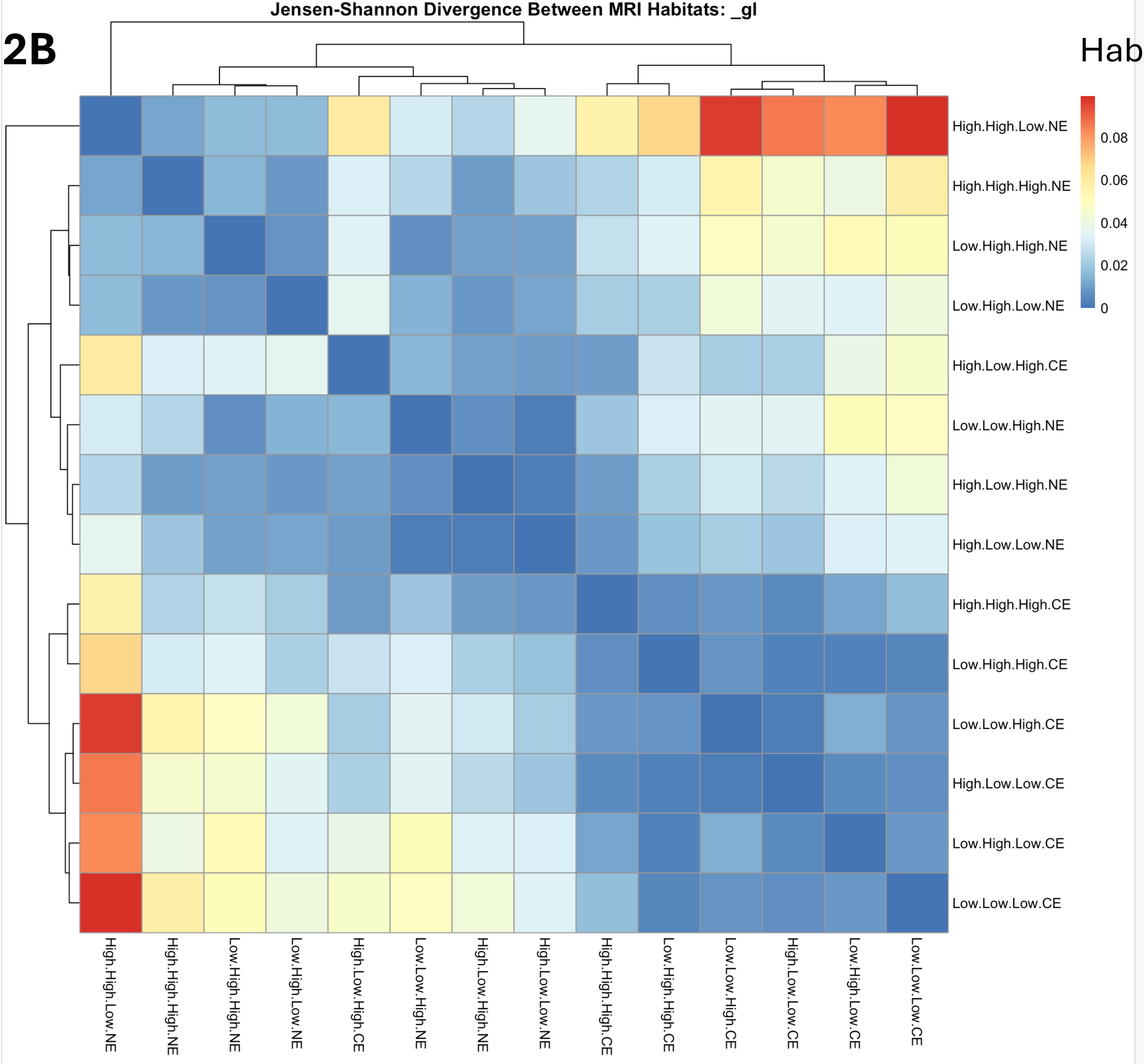


Habitat #

2
3
4
6

NE habitats with similar (lower) pseudotime and similar **glioma** cell composition

14
1
5
7

NE & CE habitats with diverse pseudotimes and similar **glioma** cell composition

10
9
13
11
12
8

CE habitats with similar (higher) pseudotime and similar **glioma** cell composition

**S12C**

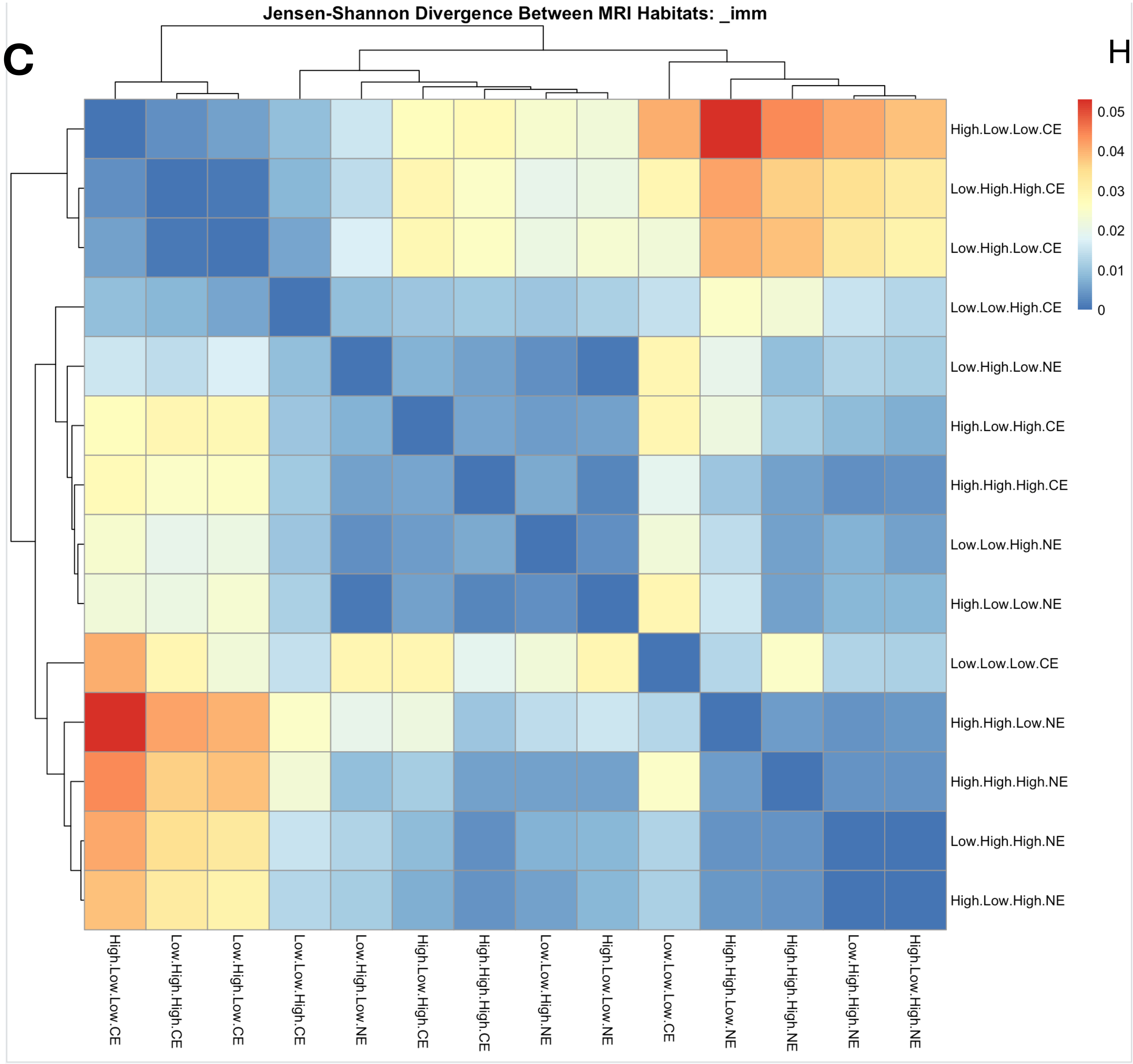


| Habitat # | |
|---|---|
| 11<br>9<br>12 | CE habitats with higher pseudotimes and similar **immune** cell composition |
| 13 | Bridging immune habitat |
| 6<br>14<br>10<br>1<br>7 | NE & CE habitats with diverse pseudotimes and similar **immune** cell composition |
| 8 | Bridging immune habitat |
| 2<br>3<br>4<br>5 | NE habitats with lower pseudotimes and similar **immune** cell composition |

**S12D**

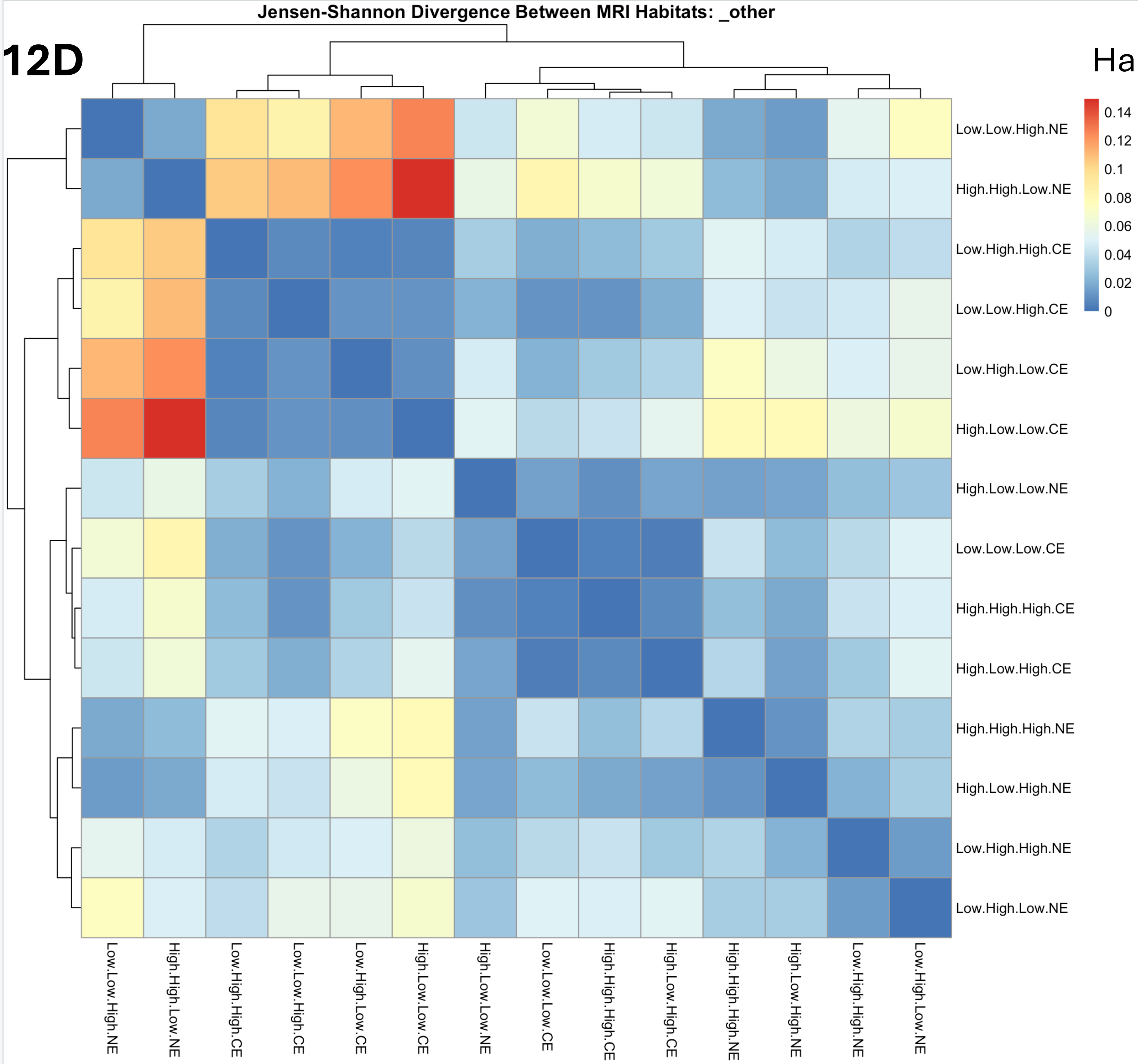


Habitat #

| | |
|---|---|
| 1<br>2 | Low pseudotime NE habitats with similar **other cell** composition |
| 9<br>13<br>12<br>11 | High pseudotime CE habitats with similar **other cell** composition |
| 7<br>8<br>10<br>14<br>3<br>5<br>4<br>6 | Moderate pseudotime CE and NE habitats with somewhat similar **other cell** compositions but also transitional between other compositions |

## S12E

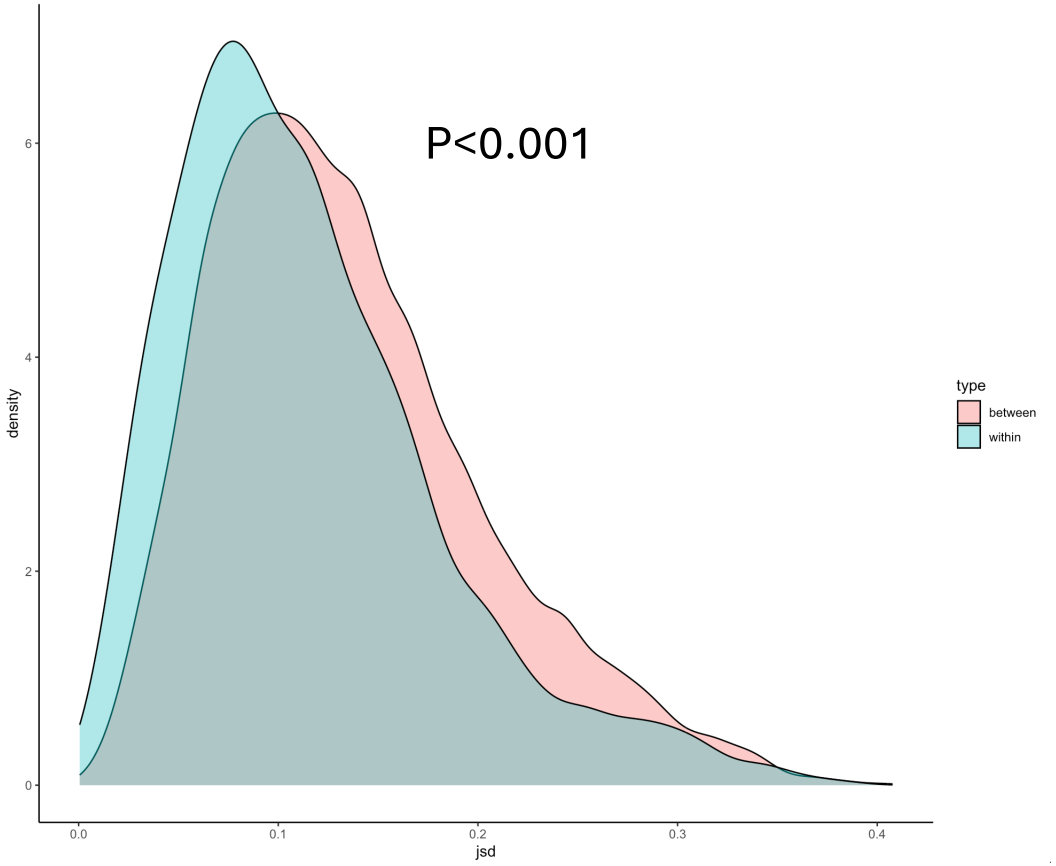


MRI Habitats are Biologically Meaningful: Comparing JSD within and between habitats shows that there is more compositional variation between habitats than within supporting the hypothesis that habitats are biologically meaningful (1000 permutations produces a mean difference of 0.021 between JSDs)

## S12F

```
>     results
              habitat within_mean between_mean      p_value
1     Low.Low.High.NE  0.11935917    0.1437799 5.824054e-02
2    High.High.Low.NE  0.08783801    0.1529639 1.045847e-09
3   High.High.High.NE  0.10315939    0.1289092 8.509811e-02
4    Low.High.High.NE  0.12411994    0.1320162 5.385243e-01
5    High.Low.High.NE  0.10978663    0.1308096 1.860012e-07
6     Low.High.Low.NE  0.13020493    0.1500202 2.462066e-02
7     High.Low.Low.NE  0.11781994    0.1307503 1.770615e-04
8      Low.Low.Low.CE  0.08620509    0.1294002 3.375157e-07
9    Low.High.High.CE  0.12881178    0.1355871 7.238213e-02
10  High.High.High.CE  0.14496822    0.1296349 3.551118e-01
11    High.Low.Low.CE  0.09480311    0.1262855 1.483336e-06
12    Low.High.Low.CE  0.11051710    0.1389701 1.058373e-32
13    Low.Low.High.CE  0.09916788    0.1230712 3.382765e-02
14   High.Low.High.CE  0.11699069    0.1397888 2.594519e-15
>
```

Some habitats are strongly distinguished by their cellular composition while others are less able to be discriminated from other habitats according to cellular composition alone. Habitats 2, 5, 6, 7, 8, 11, 12, 13, 14 are biologically distinct in terms of cellular composition from other Habitats.

Figure S12: Jensen-Shannon divergence (JSD) between MRI habitats. A. Divergence amongst all tumor cells shows a clustering of CE and NE habitats as there is more similarity between samples within those habitats rather than between habitats. There appears to be a subset of NE and CE habitats that are similar to habitats in both NE and CE habitats forming a ”bridge” between these biological states B. Divergence amongst glioma cells reveals different associations of MRI habitats in terms of similar glioma cell compositions. Similar for immune cells (C) and other non-tumor, non-immune cells (D). E. Comparing JSD within and between habitats shows that there is more compositional variation between MRI habitats than within. Performing 1000 permutations produces a mean difference between the within and between habitat JSD as 0.021 ($p<0.001$). This supports the hypothesis that MRI habitats are biologically meaningful ecological niches. F. Some habitats are strongly distinguished by their cellular composition while others are less able to be discriminated from other habitats according to cellular composition alone. Habitats 2, 5, 6, 7, 8, 11, 12, 13, 14 are biologically distinct in terms of cellular composition from other Habitats. Even when parsing JSD across MRI habitats, the pattern of more similarity within CE habitats than between CE and NE habitats was evidenced by the NE habitats clustered together and the CE habitats separately clustered together (Figure S12A). However, when we isolate the glioma cell subpopulations (Figure S12B), immune cell subpopulations (Figure S12C), and other non-tumor/non-immune cells (Figure S12D) and compute their JSD values, 3 prominent groups of habitats appeared (blue boxes). These groups mixed NE and CE regions and the associated average pseudotime for each habitat, suggesting that the spatial organization of certain tumor, immune, and other populations do not adhere to NE vs CE boundaries. We performed 1000 permutations to compare the mean difference within habitats versus across habitats to show that there was more compositional variation between MRI habitats than within (Figure S12E, $p<0.001$). Lastly, in Figure S12F, the within and between habitat JSD is shown for each habitat, identifying some habitats with strong cellular compositional distinction.

Figure S13. Continuous image features used for MRI habitats between collection sites. Linear mixed effects models with patients as a random effect did not show any significant differences between fractional anisotropy (FA, p=0.76), mean diffusivity (MD, ANOVA p=0.48), or relative cerebral blood volume (rCBV, p=0.66) values at biopsy locations collected between Barrow Neurological Institute (BNI) and Mayo Clinic Arizona (MCA). Values presented here on a log scale for clarity.

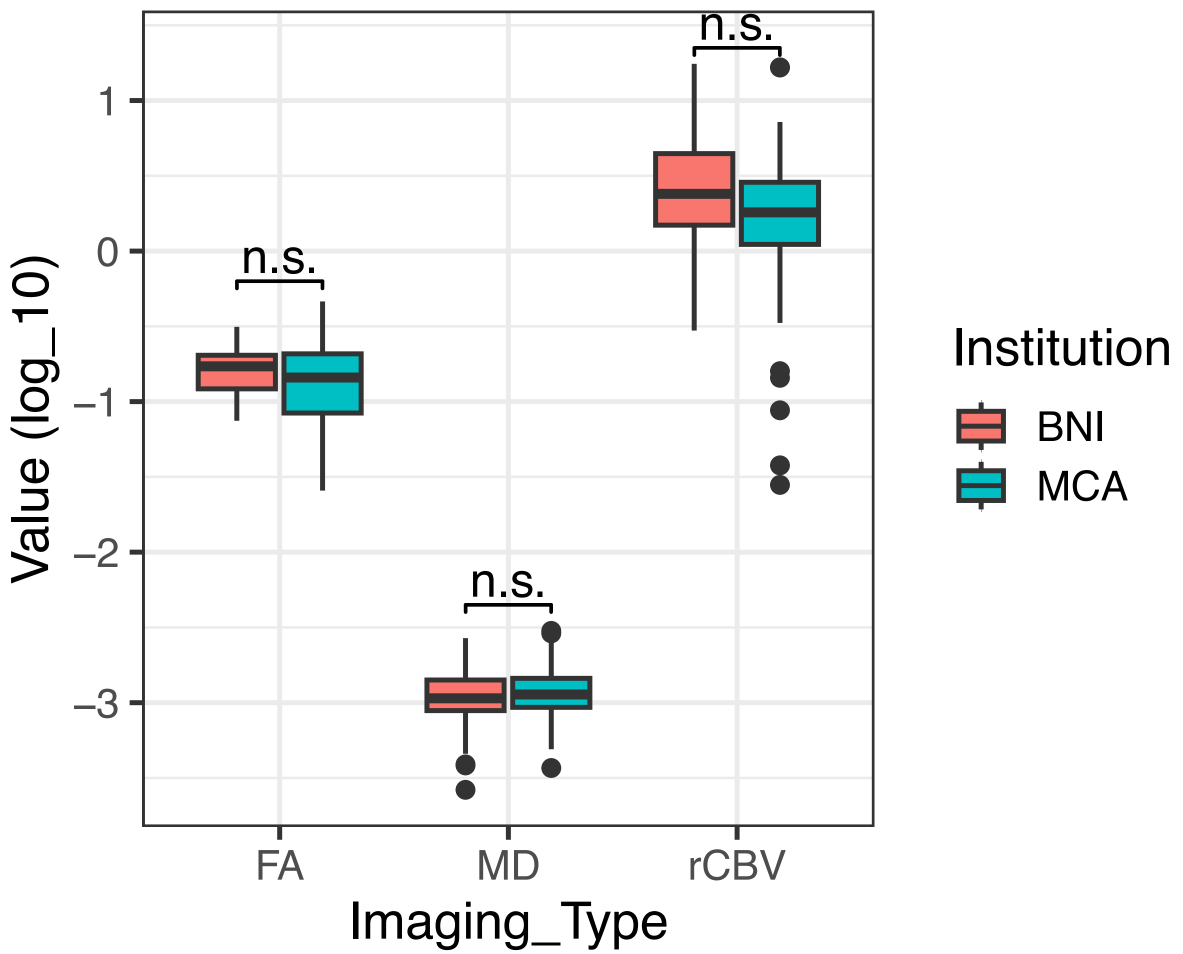

Figure S14. Leave-one-patient-out analysis. (A) Of 8016 total sample classifications, 7881 remained as their original classification (98.3%). The remaining 135 that changed in at least 1 iteration consisted of 19 unique samples from 14 unique patients (11.4%). (B) The 19 samples from 14 patients that changed habitat and within what percentage of iterations. (C) The mean absolute rank change of habitats, ranging from 0.06 to 0.61.

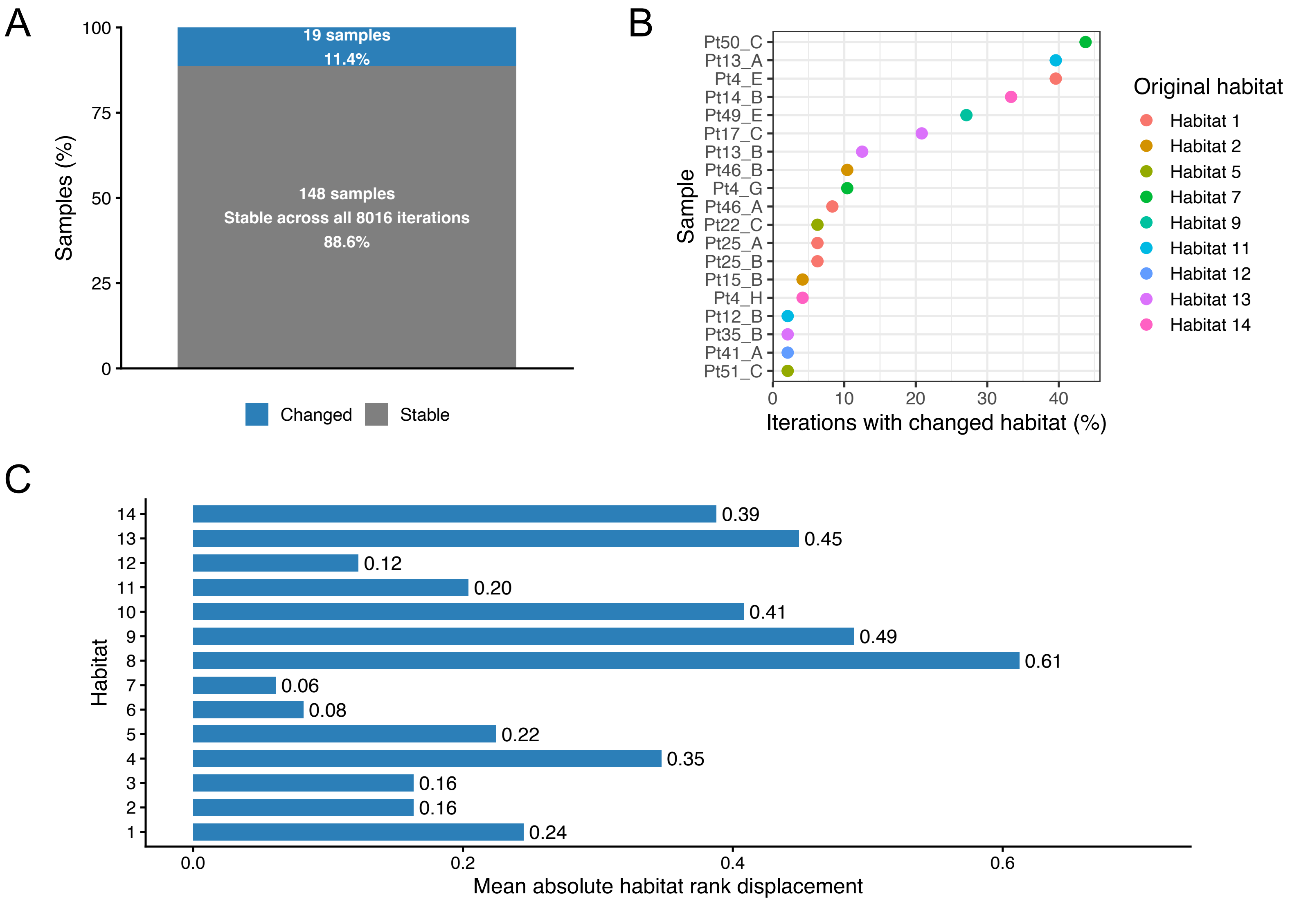

Figure S15A. Univariate cox proportional hazard models of habitat voxel counts across all 49 patients with available survival and advanced imaging

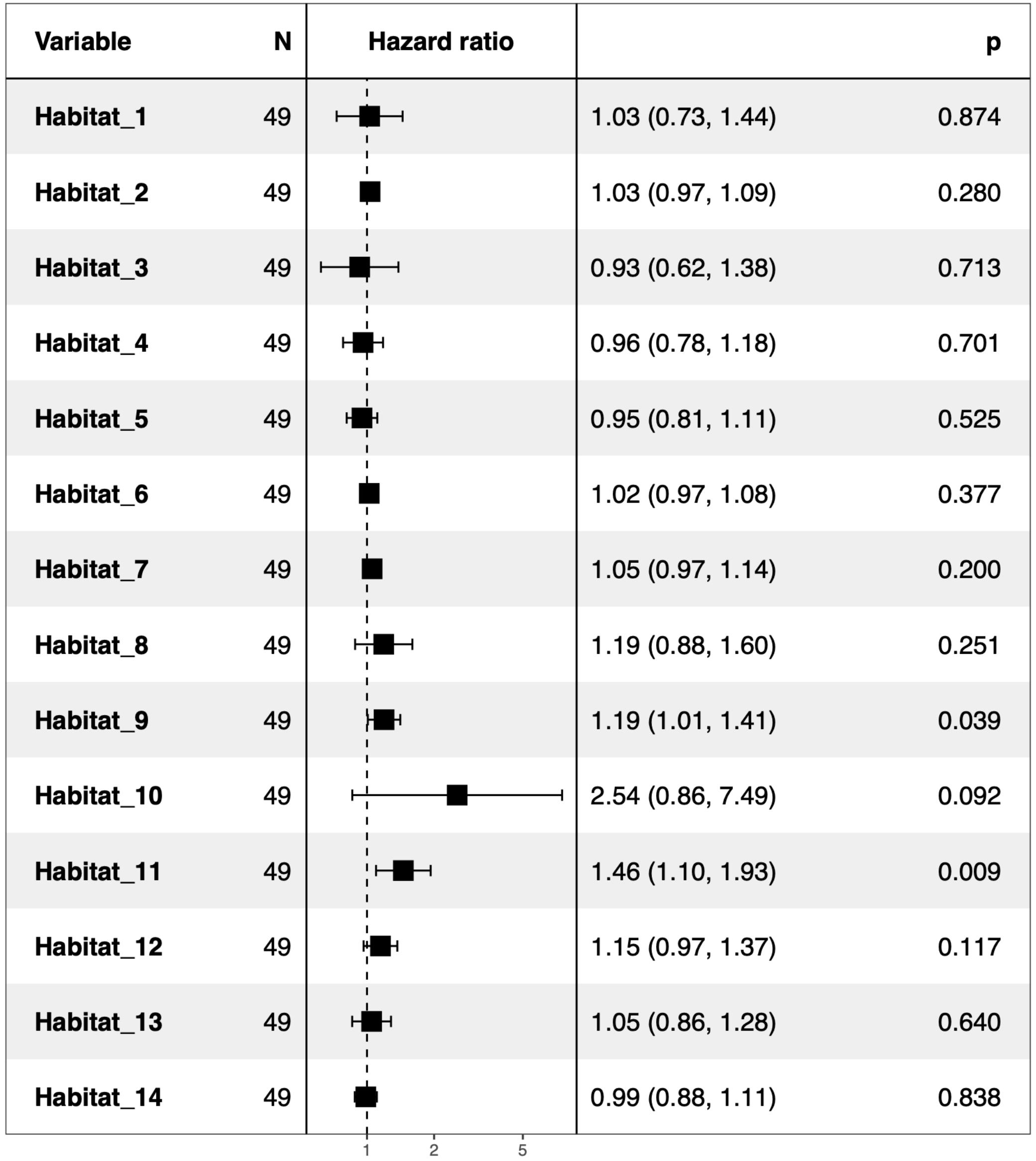

Figure S15B. Univariate cox proportional hazard models of age at diagnosis, IDH status and MGMT status

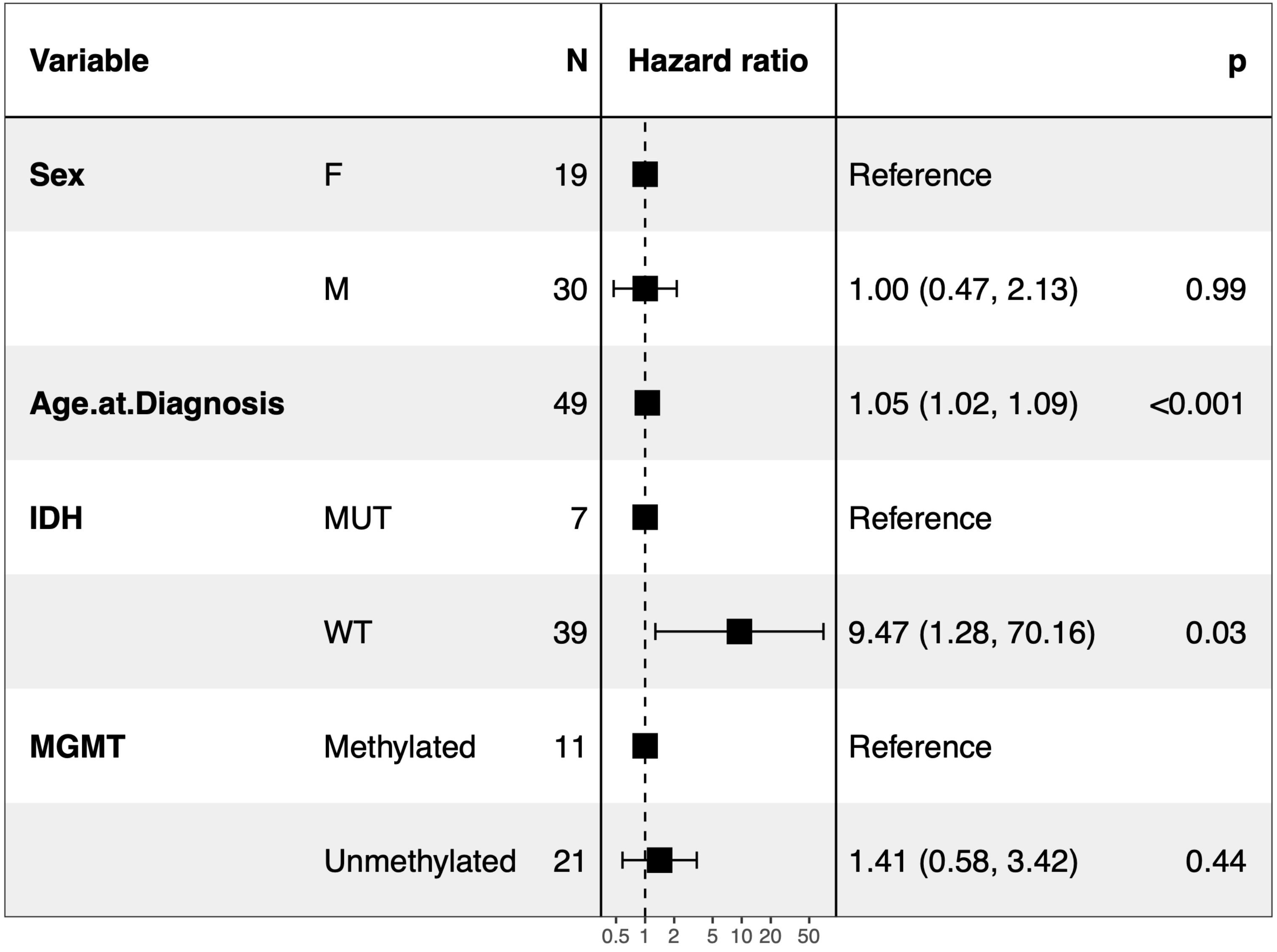

Figure S15C. Multivariate analysis of the MRI habitats with univariate p<0.1 alongside age at diagnosis

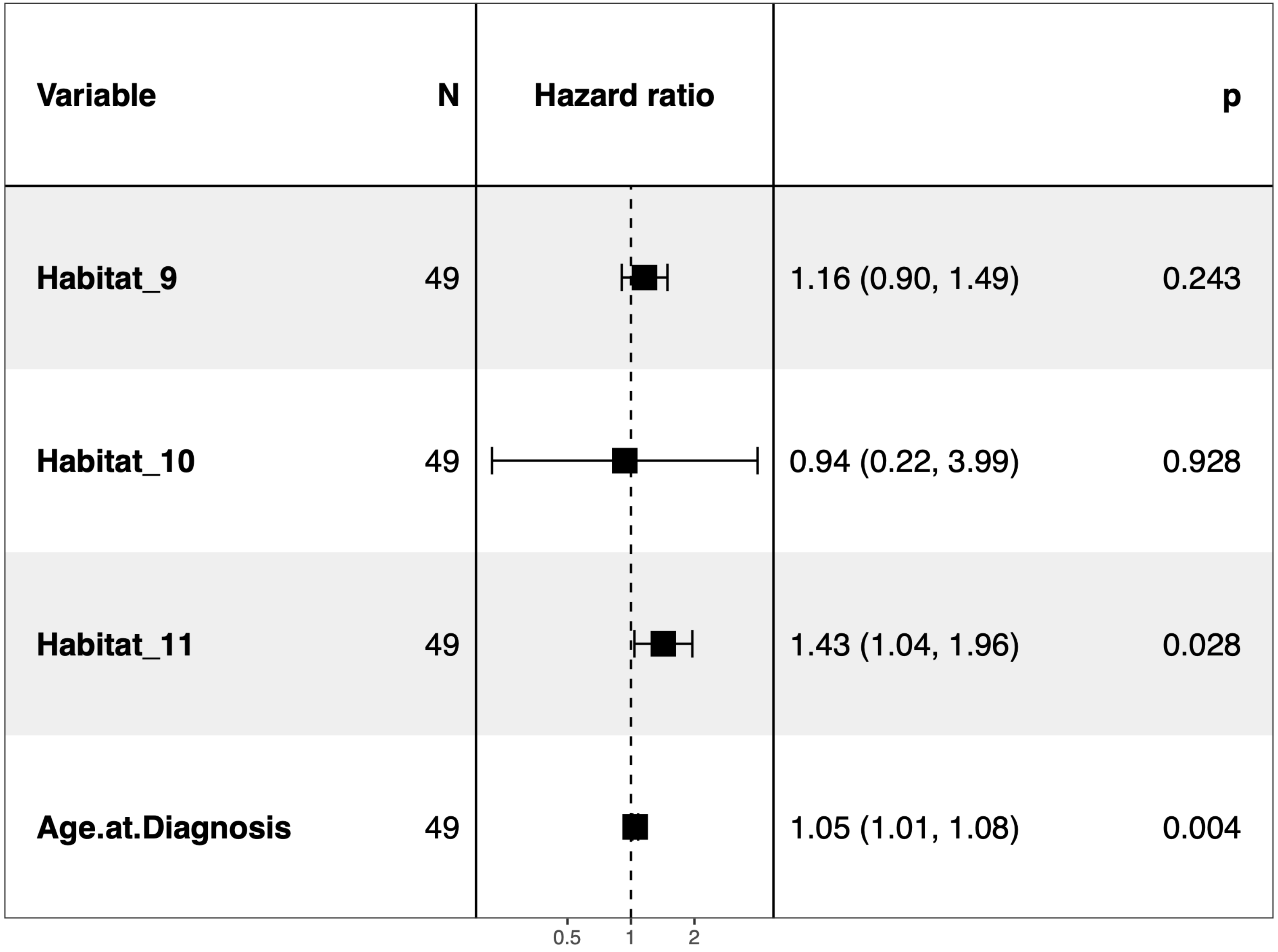

Figure S15D. Multivariate analysis of the MRI habitats with univariate $p<0.1$ alongside age at diagnosis and IDH status

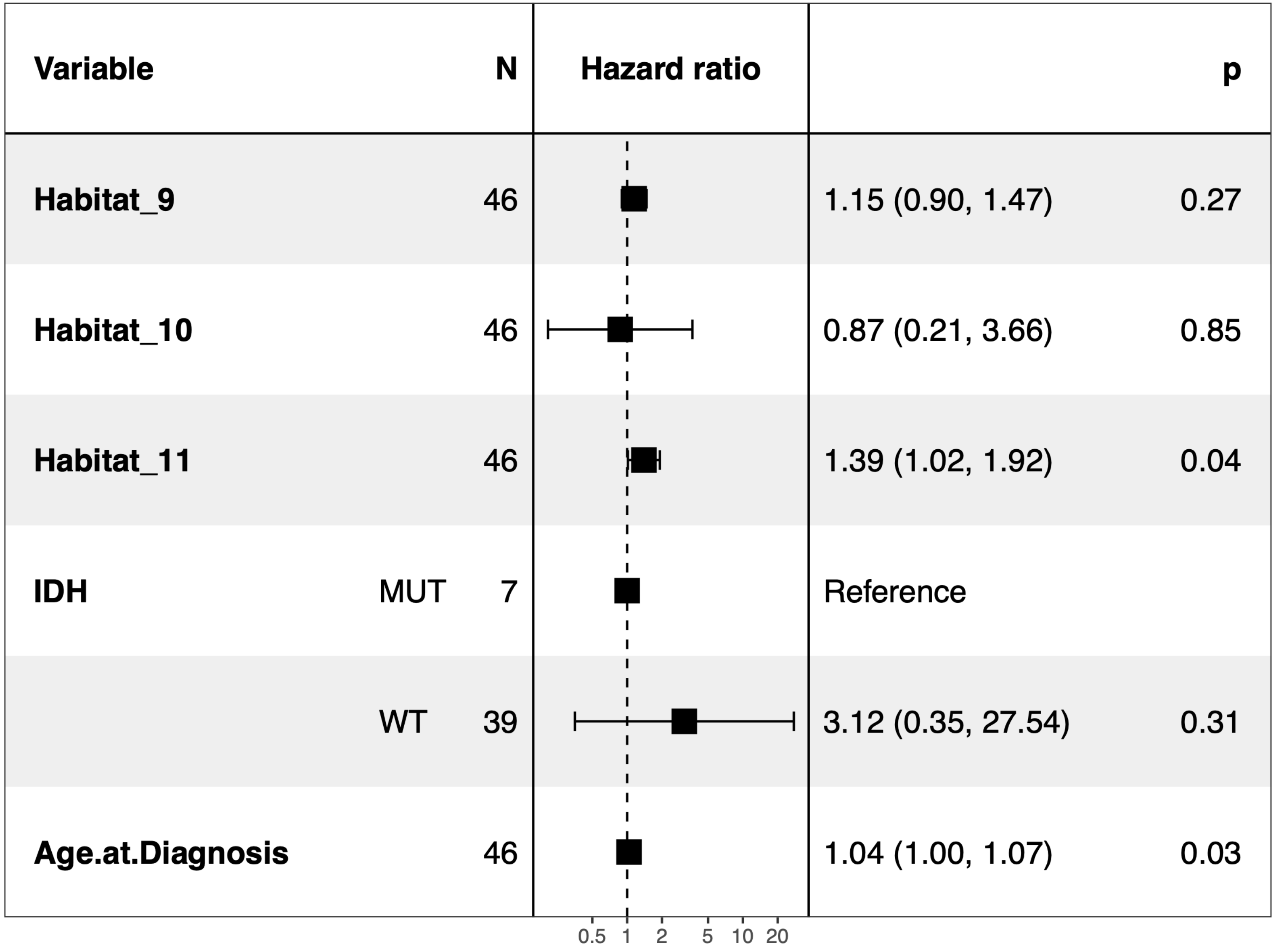

Figure S15E. Multivariate analysis of the MRI habitats with univariate p<0.1 alongside age at diagnosis and MGMT status

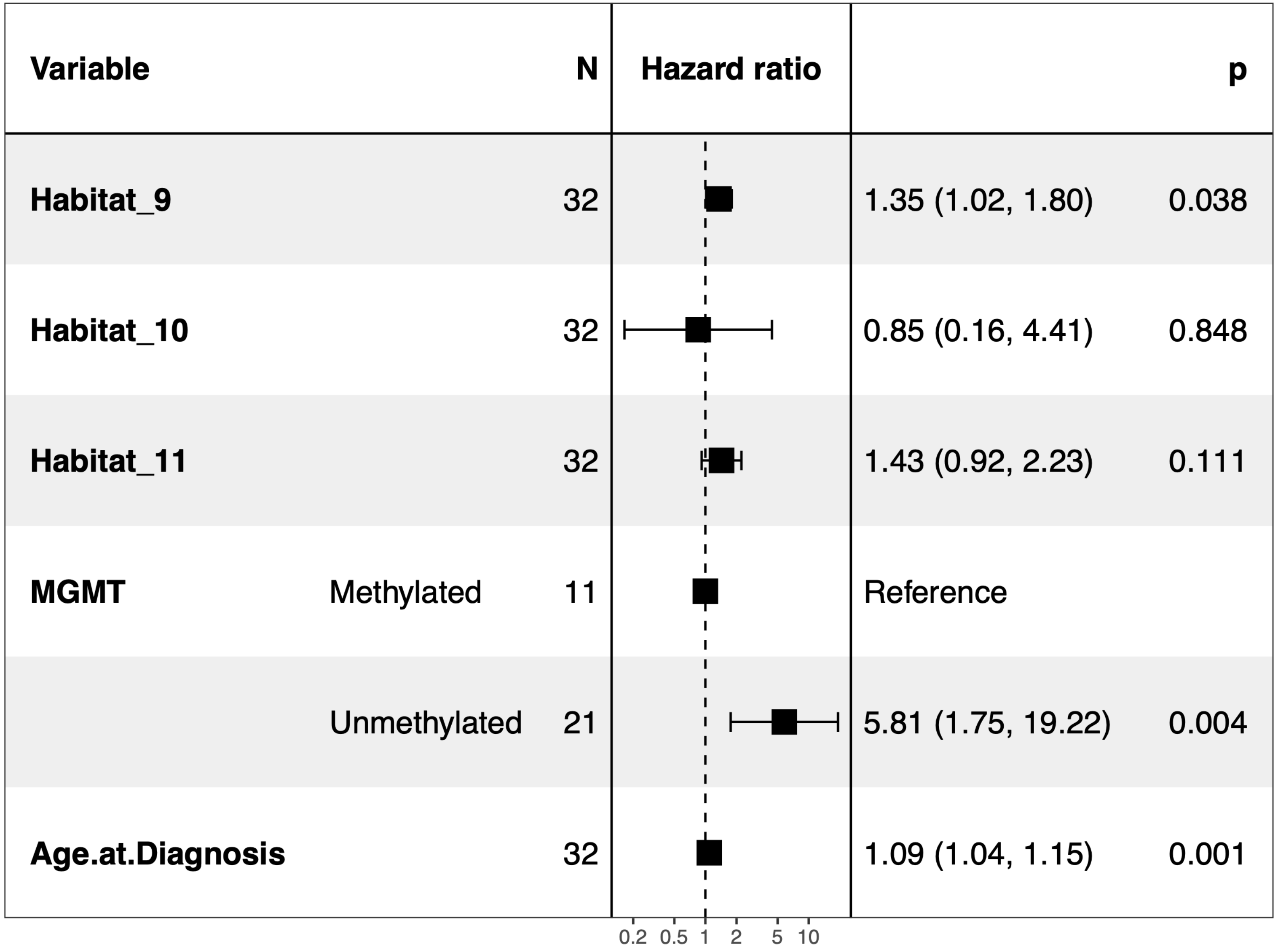

Figure S16A. FGSEA analysis of MSigDB hallmark pathways for Hallmark 9 compared to all other samples. Many cancer hallmarks are significantly upregulated in these samples (NES>0, to the right), including those associated with proliferation.

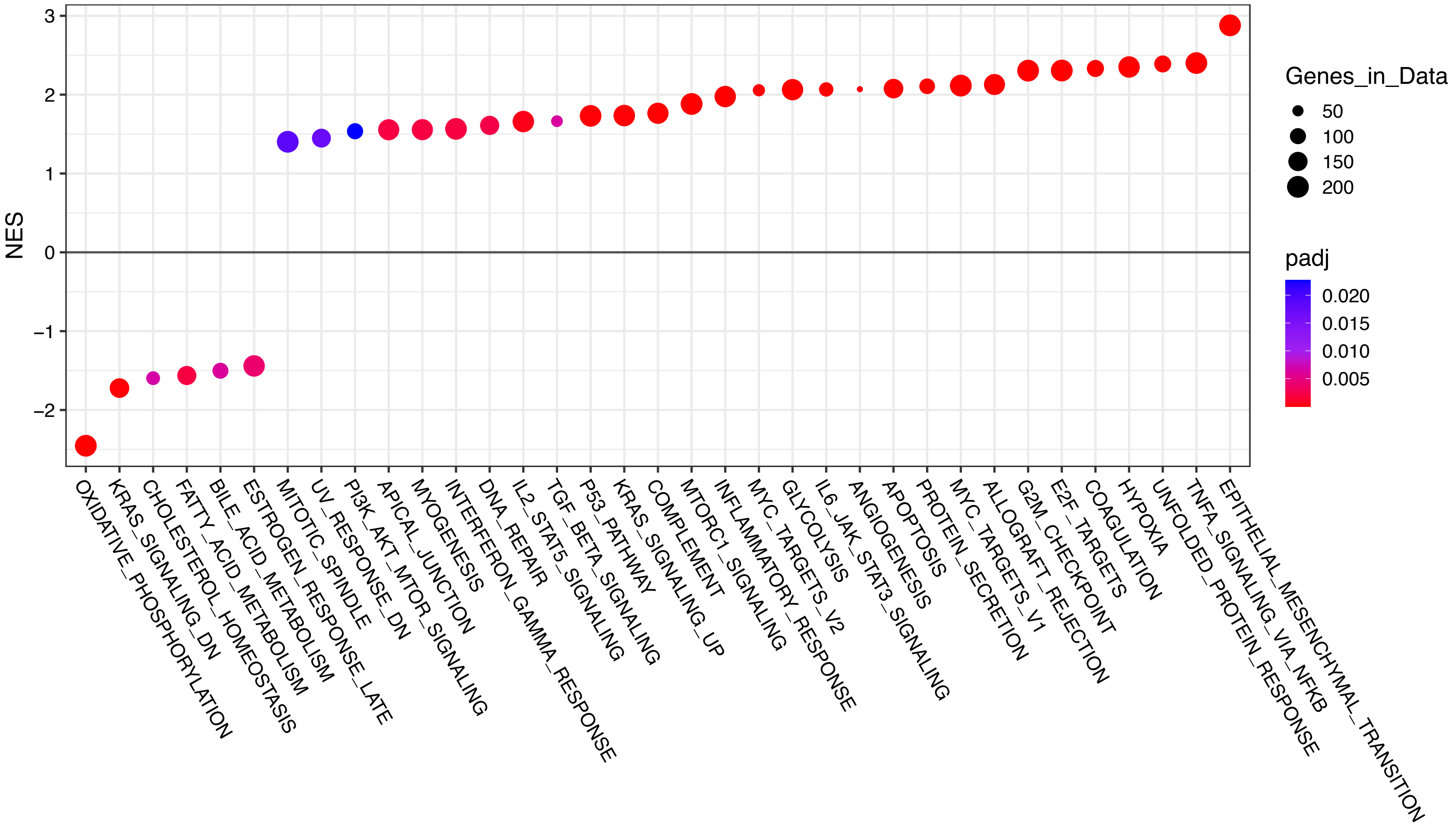

Figure S16B. FGSEA analysis of MSigDB hallmark pathways for Hallmark 11 compared to all other samples. Many cancer hallmarks are significantly upregulated in these samples (NES>0, to the right), including those associated with immune response and multiple signaling pathways.

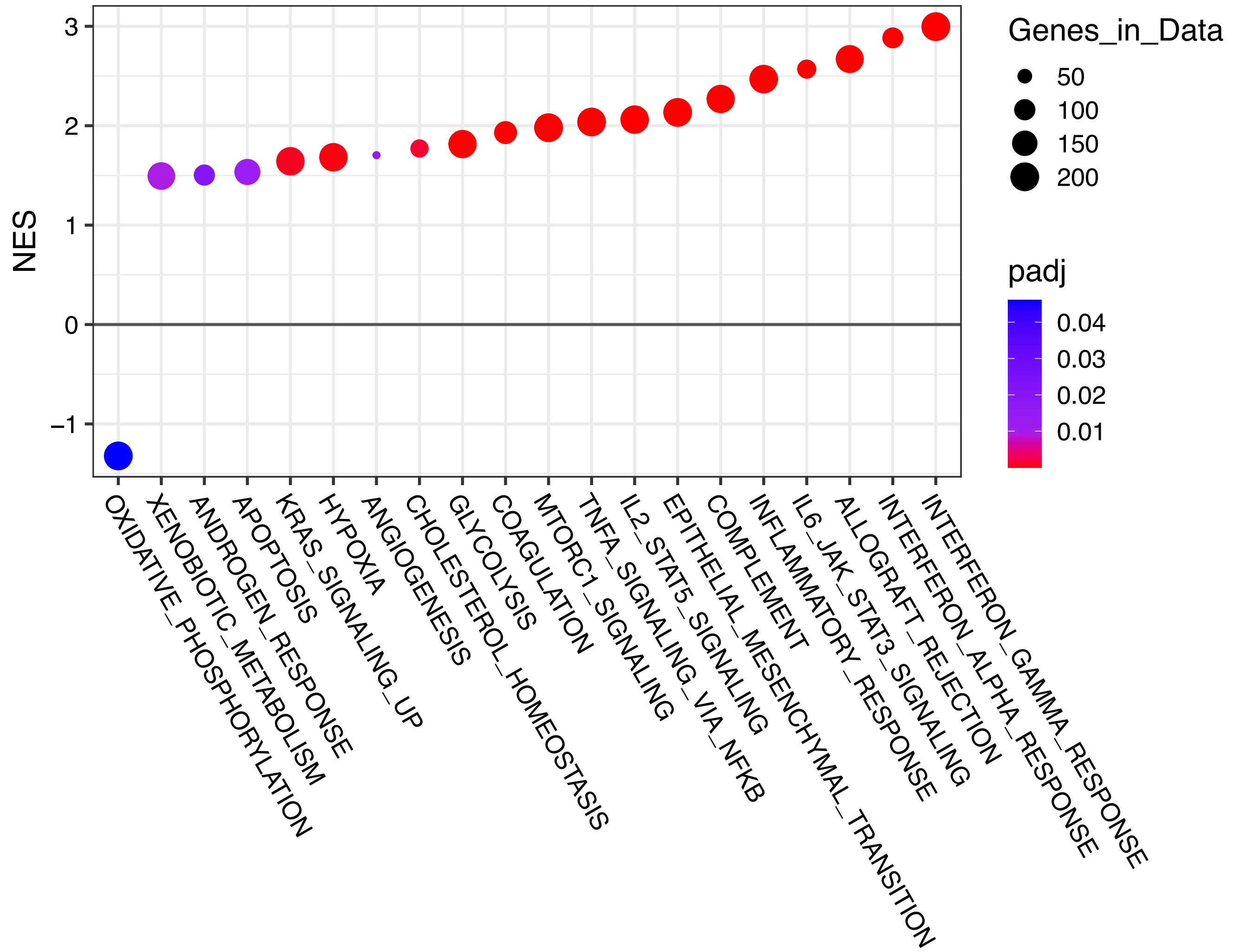

Table S1: Cell state names and descriptions

| Cell State | Description |
|---|---|
| **Malignant** | |
| **gl_Mes1** | Astrocyte-like/mesenchymal glioma - Mesenchymal / immunoreactive glioma state with astrocyte-like features |
| **gl_Mes2** | Astrocyte-like/mesenchymal glioma - Reactive astrocyte + immune-associated mesenchymal glioma state |
| **gl_Pro1** | Proliferative glioma - Cell-cycle progression module |
| **gl_Pro2** | Proliferative glioma - S-phase + DNA repair module |
| **gl_PN1** | Proneural/progenitor-like glioma - Proneural + OPC-like neural program |
| **gl_PN2** | Proneural/progenitor-like glioma - NPC-like neural program |
| **gl_sum** | Sum of all malignant states |
| **Immune** | |
| **Tcell** | Adaptive immune effector/infiltration module |
| **Myel1** | Myeloid inflammatory/metabolic stress module |
| **mgTAM** | Microglia-derived tumor-associated macrophage state |
| **moTAM** | Monocyte-derived tumor-associated macrophage immunosuppressive state |
| **prTAM** | Proliferative perivascular remodeling tumor-associated macrophage state |
| **immune_sum** | Sum of all immune states |
| **Other** | |
| **Neuron** | Neuronal program |
| **OPC** | Oligodendrocyte progenitor cell lineage |
| **Oligodendrocyte** | Mature oligodendrocyte lineage |
| **Endothelial** | Endothelial cell |
| **Ast1** | Protoplasmic astrocyte-like, interface/homeostatic state |
| **Ast2** | Reactive astrocyte with plasticity/tumor–glial hybridization |
| **Ast3** | Inflammatory astrocyte module |
| **ast_sum** | Sum of astrocyte populations Ast1, Ast2 and Ast3 |
| **other_sum** | Sum of all other states excluding ast_sum |

Table S2: Patient level statistics

| **Characteristic** | **N = 58**[1] |
|---|---|
| Age (years) | 61 (21-84) |
| Sex | |
| F | 22 / 58 (38%) |
| M | 36 / 58 (62%) |
| IDH | |
| Mut | 8 / 54 (15%) |
| Wt | 46 / 54 (85%) |
| Unknown | 4 |
| MGMT | |
| Methylated | 11 / 36 (31%) |
| Unmethylated | 25 / 36 (69%) |
| Unknown | 22 |

[1]Median (Min-Max); n / N (%)

#### Table S3: Sample level statistics

| Characteristic | N = 202[1] |
|---|---|
| Sex | |
| F | 80 / 202 (40%) |
| M | 122 / 202 (60%) |
| IDH | |
| Mut | 21 / 189 (11%) |
| Wt | 168 / 189 (89%) |
| Unknown | 13 |
| MGMT | |
| Methylated | 45 / 127 (35%) |
| Unmethylated | 82 / 127 (65%) |
| Unknown | 75 |
| Status | |
| P | 153 / 202 (76%) |
| R | 49 / 202 (24%) |
| Age.at.Diagnosis | 63 (21-84) |
| Enhancement | |
| CE | 123 / 195 (63%) |
| NE | 72 / 195 (37%) |
| Unknown | 7 |
| MD ($10^{-3}$ $mm^2/s$) | 1.12 (0.26-3.00) |
| Unknown | 31 |
| FA | 0.16 (0.03-0.46) |
| Unknown | 31 |
| rCBV | 2.00 (0.00-17.53) |
| Unknown | 25 |
| State | |
| 1 | 69 / 202 (34%) |
| 3 | 65 / 202 (32%) |
| 4 | 44 / 202 (22%) |
| 5 | 24 / 202 (12%) |
| Pseudotime | 38 (0-59) |

[1]n / N (%); Median (Min-Max)

Table S4: Monocle states with their biological labels, dominant CIBERSORTx populations, and associated imaging regions.

| Monocle State | Biological Label | Dominant Cellular Populations | Dominant Enhancing Status |
|---|---|---|---|
| **State 1** | Mesenchymal Immune/Inflammatory | gl_Mes1, gl_Mes2, moTAM, Endothelial, Ast3, Myel1 | Contrast Enhancing |
| **State 2** | NA | NA | NA |
| **State 3** | Invaded Brain | Neuron, Ast1, Oligodendrocyte | Non-Enhancing |
| **State 4** | Proliferative | gl_Pro1, gl_Pro2, gl_PN2, prTAMs, OPC | Contrast Enhancing |
| **State 5** | IDH Mutant Dominant | gl_PN1, mgTAM, Myel1, Oligodendrocyte | Contrast Enhancing/Non-Enhancing Mix |
