## Supplementary Methods for "Mapping multiregional image-localized biopsies to MRI habitats reveals biologically significant glioma tissue states and patterns of cellular subpopulations"

Tissue samples

These samples were collected from Barrow Neurological Institute (n=51) and Mayo Clinic (n=151). We obtained three-dimensional coordinates for these samples and also classified them more broadly into contrast-enhancing (CE; from regions with T1Gd abnormalities) and non-enhancing (NE; from other regions, enhancing on T2/FLAIR MRI and in normal-appearing brain) as these distinctions are central to surgical planning and the radiological assessment of treatment responsiveness. Biopsy locations on registered images were verified by three individuals (L.C., J.C.U., K.W.S.) as were CE/NE calls (L.S.H., K.W.S., K.M.B). Further methods for the image-localized sample collection protocol are presented here[60]. The tissue was flash-frozen, and the Illumina TruSeq v2 RNAseq kit was used to prepare sequencing libraries. An Illumina HiSeq 4000 sequencer obtained paired-end reads, and FASTQ files were aligned to a reference genome (GRCh38.p37). Read counts were compiled using htseq-count, and batch effects were corrected with ComBat-Seq[66]. The sex of each patient in this work was self-reported. *TCGA Biopsy Data:* 525 patients (205F, 320M) with GBM underwent tissue characterization as a part of The Cancer Genome Atlas (TCGA) effort. Microarray data from these samples were accessed using UCSC Xena[67].

MRI acquisition, image localization and segmentation

Pre-contrast T1-weighted (T1W), T2-weighted (T2W), fluid-attenuated inversion recovery (FLAIR), diffusion tensor imaging (DTI), and perfusion weighted imaging (PWI) sequences were acquired at 3T field strength. Acquisition parameters were as follows: T1W (TR/TE=600/12ms, matrix=320x240, FOV=24cm, thickness=2mm), T2W (TR/TE=4000/76ms, matrix=384×384; FOV=22cm; thickness=3mm), FLAIR (TI/TR/TE=2600/10030/135ms, matrix=320x320, FOV=24cm, thickness=2mm), and DTI (TR/TE = 6700/78ms, matrix=150x150, FOV=22cm, thickness=2.6mm, b-value=1000 s/mm$^2$, number of directions = 30). Dual flip angle PWI was performed with two acquisitions at 30° and 60° with a dose of gadolinium-based contrast agent given with each acquisition (TR/TE=1500/20ms, matrix = 150x150, FOV=22cm, thickness=2.6mm). Post-contrast T1W images (T1Gd) were acquired following PWI. Using previously described methods, MRI coordinates for image-localized biopsies were captured, images were normalized, co-registered and abnormalities segmented[60,68,69]. Advanced imaging mean values were determined using denoising, N4 bias field correction, z-score normalization in SimpleITK, and PyRadiomics for final feature extraction.

Deconvolving cell populations from RNA sequencing

While many have employed single-cell RNA sequencing as a means of characterizing intratumoral heterogeneity, this approach notoriously undersamples non-neoplastic populations (e.g., neurons)[70]. Single nucleus RNAseq (snRNAseq) circumvents this limitation by more comprehensively sampling malignant, immune, and other cells. Malignant cell populations included mesenchymal/immunoreactive (gl_Mes1 and gl_Mes2), proliferative (gl_Pro1 and gl_Pro2), and neural/oligodendrocyte precursors (gl_PN1 and gl_PN2). Immune cells were characterized as T-cells, myeloid cells present at baseline (Myel1), proliferative tumor-associated macrophages (prTAM), monocyte-derived cells (moTAM), and microglia-derived cells (mgTAM). Other populations were characterized as neurons, endothelial cells, oligodendrocytes, oligodendrocyte precursor cells (OPCs), and three subtypes of astrocytes (Ast1, protoplasmic; Ast2 and Ast3, reactive) (Table S1)

Before running the algorithm, sex-linked genes of *XIST*, *DDX3*, *EIF1AY*, *KDM5D*, *NLGN4Y*, *ZFY*, *RPS4Y1*, *TMSB4Y*, *USP9Y*, and *UTY* were removed in the bulk RNA-Seq dataset. These were removed to avoid the CIBERSORTx algorithm biasing cellular abundance towards sex-linked genes. Signature matrices were then generated for the bulk RNAseq samples to estimate the abundance of all eighteen cell types in our image-localized samples. We downsampled the snRNAseq data to 100 cells of each type three times, and ran each of these three times to increase the robustness of our output to algorithm variability. We present means across these nine runs for each sample and cell type abundance throughout. Further analysis of robustness within and between down samples can be found in the supplement (Figure S1). This analysis was carried out in relative mode. Gene names were aligned using the GeneSymbolThesarus package between the image-localized and snRNA datasets[71,72]. S-mode batch correction was applied to account for the application of a signature derived from UMI-based snRNAseq to bulk RNAseq data. All other CIBERSORTx parameters were kept as their default.

Trajectory inference and pseudotime ordering of TCGA microarray and image-localized RNAseq samples

Monocle reduces data dimensionality using independent component analysis (ICA).[40] A manifold learning algorithm (DDRTree) uses reverse graph embedding to identify potential backbones of the trajectory and orders samples along the longest path. Each sample is assigned to a branch on the tree, and pseudotime is calculated using the geodesic distance to the path's starting point. Here we used TCGA microarray and our biopsy RNAseq as inputs to visualize the natural transcriptional organization of HGG. States for our image-localized biopsy data are presented as assigned by Monocle.

Differential Gene Expression and Gene Set Enrichment Analysis

Differential expression was used to assess which gene sets were significantly over/underexpressed in a variety of data stratifications, we used the "limma" package in R with a patient block design to account for multiple samples per patient[34]. Lowly-expressed genes were excluded using voom, and a ranked list of genes was then created. The "fgsea" package was then used to perform gene set enrichment analysis (GSEA), with the hallmarks pathways from the MSigDB database as the reference[35]. These results were used to assess the statistical significance of the hallmark signatures between varying metadata groupings of the data, with the Benjamani-Hochberg used as a p-value adjustment for multiple comparisons. Single sample GSEA was used to determine the pathway expression of each sample using the "gsva" package in R[36]. HGNCHelper was used for alignment between the image-localized and TCGA datasets[37]. Up to the top 100 upregulated and downregulated genes associated with each Monocle state of the 202-sample dataset were used alongside single-sample GSEA (ssGSEA) to create state scores for each TCGA microarray sample. The state with the highest score was then assigned as the predicted state for each sample across the TCGA microarray dataset.

*MGMT* Methylation Inference

Due to the lack of available *MGMT* methylation data in the image-localized cohort (75/202 samples without values), we used an optimized *MGMT* expression threshold to infer methylation status. This was calculated using a receiver operator curve in the "pROC" R package[73] on the averaged *MGMT* expression per patient for CE samples only, to more closely replicate the clinical process for observing methylation status. This simple threshold model had an AUC of 0.81, with the optimal counts threshold of 203.5 leading to 0.68 sensitivity and 0.78 specificity. These inferred *MGMT* methylation values are presented only to visualize trends and are excluded within statistical analyses.

Statistical analyses

Since the TCGA dataset only contained one sample per patient, we used standard methodology for statistical analyses applied to this dataset as the independence of observations assumption can be met (chi-square, pearson correlations, log-rank). We did not assume independence for our image-localized biopsy cohort as most patients had multiple samples collected during surgery, thus linear mixed-effects models were used to determine the significance of differences between population abundances within this dataset, with box plots employed for data visualization. Patient IDs were included as a random effect in each test within the lmer functions in the "lmerTest" R package[39], with Satterthwaite's method used for comparisons of 2 groups, and anova tests implemented for comparisons with 3+ groups. A Benjamani-Hochberg correction was used when adjusting for multiple comparisons.
